# Activity-dependent structural plasticity of myelinated axons tunes their excitability

**DOI:** 10.64898/2026.08.07.743576

**Authors:** David J. Bennett, Vladimir Rancic, Temitayo Ayantayo, Zhenxiang Zhao, Mohammed Almokdad, Claire F. Meehan, Keith K. Fenrich, Ana Lucas-Osma, Krishnapriya Hari

**Affiliations:** Neuroscience and Mental Health Institute, University of Alberta, Edmonton, Alberta, Canada; Faculty of Rehabilitation Medicine, University of Alberta, Edmonton, Alberta, Canada; Department of Neuroscience, University of Copenhagen, Panum Institute, Blegdamsvej 3, 2200 Copenhagen N, Denmark

**Keywords:** axonal plasticity, dorsal column, node of Ranvier, Kv1, Na/K-ATPase, glia, primary afferent depolarization, axon excitability, epidural stimulation

## Abstract

While myelinated axons are classically viewed as fixed cables, recent evidence suggests that activity can rapidly alter their geometry. Here we examined how this occurs and its functional implications, by using two-photon imaging, excitability testing and axial conductance measurements in adult mouse spinal cords. We found that focal subthreshold stimulation thins the internode to about 70% of its diameter within seconds, sparing the node and paranode, while neighbouring glia swell to fill the vacated space. The thinned axon becomes markedly more excitable for hours, by the amount cable theory predicts from its shortened length constant. Thinning requires Kv1 channels under the myelin, Na/K-ATPase, bicarbonate and a local alkaline shift, and is reproduced and occluded by axonal GABA_A_ receptor activation. Clinically used charge-balanced stimulation protocols drive the same change. Activity-dependent axon thinning therefore gives axons a structural memory of recent activity, and may underlie the lasting benefits of spinal cord stimulation.

## INTRODUCTION

Myelinated axons conduct with speed and fidelity because their geometry is precisely matched to the distribution of ion channels at the nodes of Ranvier. The axon diameter, in particular, sets the conduction velocity and current required to bring a node to threshold, and is conventionally treated as a fixed property of the mature fibre^1,2^. However, increasing evidence indicates that axon activity can change axonal structure^3–5^. For example, prolonged activation of many myelinated axons simultaneously can cause the paranodal regions to expand and the internodal regions to shrink for seconds to hours afterwards, a reversible osmotic change that involves the Na/K-ATPase pump^4^, but does not necessarily require axon spikes^5^. Whether such structural plasticity alters axon function and has a physiological role is unclear. If so, it may provide both a novel kind of memory of bulk axon activity, and a mechanism that might explain the bewildering range of brain stimulation paradigms that provide clinical benefits, including stimulation near or below threshold where a local non-spiking mechanism must come into play. This question is sharpened by observations from stimulating the intact spinal cord, where a brief subthreshold polarization of dorsal column axons produces a dramatic increase in their excitability that outlasts the stimulus by many hours^6,7^. Furthermore, epidural stimulation or transcutaneous spinal cord stimulation that engages these axons^8,9^ is used clinically to modify sensory and motor function^10–12^, sometimes long after the current is switched off^13,14^. The cellular basis of these lasting changes is not known, with it being especially unclear how a subthreshold stimulus drives a persistent change in axons. Here we combine live imaging, and axon excitability and conductance testing to show that focal subthreshold stimulation of dorsal column axons drives a rapid, lasting internode thinning that increases excitability through a purely geometric, cable-theoretic mechanism, and that the thinning is initiated by potassium currents under the myelin and Na/K-ATPase, and accompanied by glial swelling. Importantly, the same structural plasticity is engaged physiologically by axonal GABA_A_ receptors and can become pathological.

## RESULTS

### Focal subthreshold stimulation thins the axon internode region and raises the axon excitability

To examine how an activity-dependent structural change in axons influences their function, sensory axons were labelled with a peripheral AAV9 viral vector injection and the whole adult sacral spinal cord was extracted in vitro for two-photon imaging of these axons in the dorsal columns. We focally activated single nodes of dorsal column axons with a weak depolarizing subthreshold current lasting 1 to 3 s, delivered through a tungsten electrode positioned about 50 μm from the spinal cord surface to mimic epidural stimulation. This produced a rapid and long-lasting thinning of large myelinated axons to about 70% of their initial diameter, a 50% reduction in volume, starting within 2 s of the stimulus, peaking at 10.48 ± 2.19 s and lasting > 2 h over which we imaged (Fig 1A). We focused on group I and II axons for analysis, though smaller axons also shrank (Fig 1A). Immunolabelling the nodal region for NaV, caspr and Kv1.2 showed that the node and paranode remained unchanged while the internode shrank (Fig 1B; diameter measurements of axon, not including myelin). This was also seen in live imaging where nodes of the largest axons are evident from their characteristic constrictions prior to stimulation (Fig 1A). Only the internode adjacent to the stimulated node shrank, and not internodes of more distal nodes (Fig 1A) or laterally far from the electrode (top of Fig 1A), consistent with a non-propagated, subthreshold activation confined to one node. As the axon shrank, neighbouring glia swelled to fill the space it vacated, so that adjacent axons remained roughly in place rather than being drawn closer together (Fig 1A; black gaps wider). Immunolabelling astrocytes with GFAP showed that their periaxonal processes became swollen (Fig 1C; see also irregular glial swelling bending axons at arrows in Fig 1A). The myelin also swelled with irregular bulges, and stayed closely apposed to the axon without a gap opening between them, as seen by labelling the non-compacted myelin with CNPase (Fig 1D), presumably because the fluid-filled inner tongue of the myelin swelled to take up the difference. In the in vivo urethane anesthetized mouse axons shrank similarly with the same dorsal column stimulation, to mimic epidural stimulation (n = 12 of 12 tested, not shown). Furthermore, axon thinning also occurred for trains of short pulses (1 min, 100 Hz; Fig 1A).

**Figure 1.**
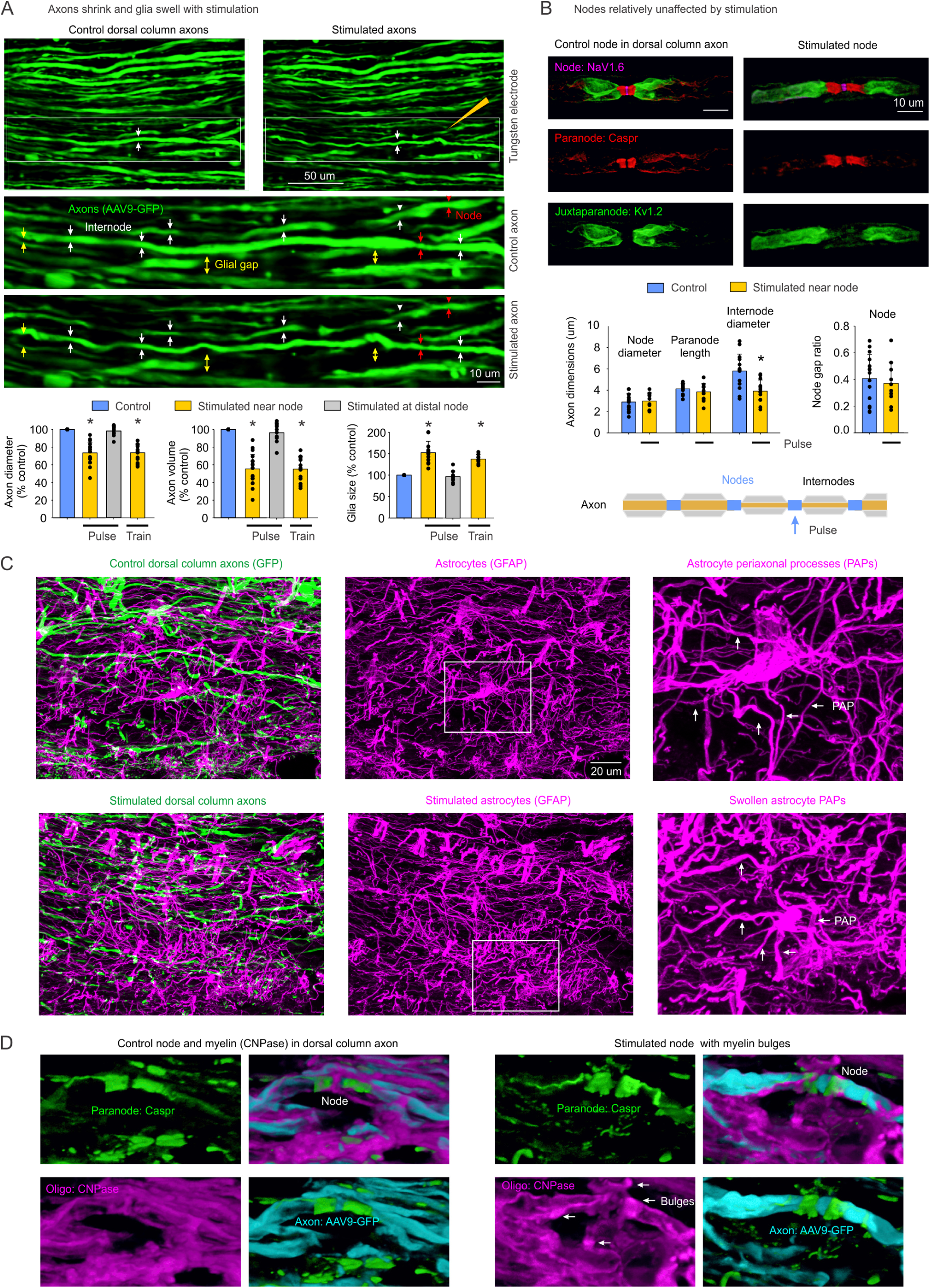
Stimulation thins the internode and swells the surrounding glia. A, Two photon imaging of live dorsal column axons labelled with AAV9-GFP, before and after focal subthreshold stimulation with a large tungsten electrode, with either a 3 s pulse (cathodal current, 15 to 20 μA, < Threshold, T = 25 μA; top images and group data) or a 1 min train (100 Hz, 20 μA, 0.4 ms pulses; group data shown); 10 μm z stack. Stimulation at location indicated (gold large electrode symbol, but tip 55 μm wide), and influenced nearby axons most. Insets show the expanded axon, with axon shrinking and glial swelling indicated with arrows. Group data show a reduced axon diameter and volume and an increased glia size near the stimulated node for both the pulse and the train (percent of control; over 200 μm range laterally), with internodes at distal nodes unchanged. Net glia size estimated from gap between axons (black). * significant change, p < 0.05, n = 12 mice each. B, Immunolabelling in control and stimulated axons, showing the node (NaV), paranode (Caspr) and juxtaparanode (Kv1.2), with group data showing that the internode diameter is reduced while the paranode length, node diameter and node gap ratio (node length/diameter) are unchanged. * significant change, p < 0.05, n = 13 to 15 each. C, Astrocytes (GFAP) around control and stimulated axons, showing increased GFAP volume and thickened periaxonal astrocyte processes (PAPs, arrows) after stimulation. D, Non-compact myelin (CNPase) at control and stimulated nodes, showing that stimulation produces myelin bulges (arrows). Similar results observed for n = 11 paired control/stimulation experiments in C and D, where axons compared from the left (stimulated) and right (unstimulated) dorsal columns.

The same stimulation dramatically increased the excitability of the axons over a similar time course (7.78 ± 3.67 s, n = 25 mice, not significantly different from shrinking, p > 0.05). Tested with a brief 0.1 ms pulse applied through the same tungsten electrode, the stimulated axons showed 3-fold increases in the compound action potential (CAP) that was recorded propagating out the dorsal roots to a recording wire mounted in grease (324.9 ± 93.98% of control, n = 25; Fig 2), matching the marked excitability increases reported for comparable stimulation in vivo^6^. When the dorsal column stimulation was applied close to the recording site, within a few internodes, a large local depolarization associated with the increased excitability was observed (Fig 2B, detailed below), but this depolarization was not propagated, as more distant stimulation caused only an increased CAP (Fig 2C), again consistent with a local subthreshold action on nodes.

**Figure 2.**
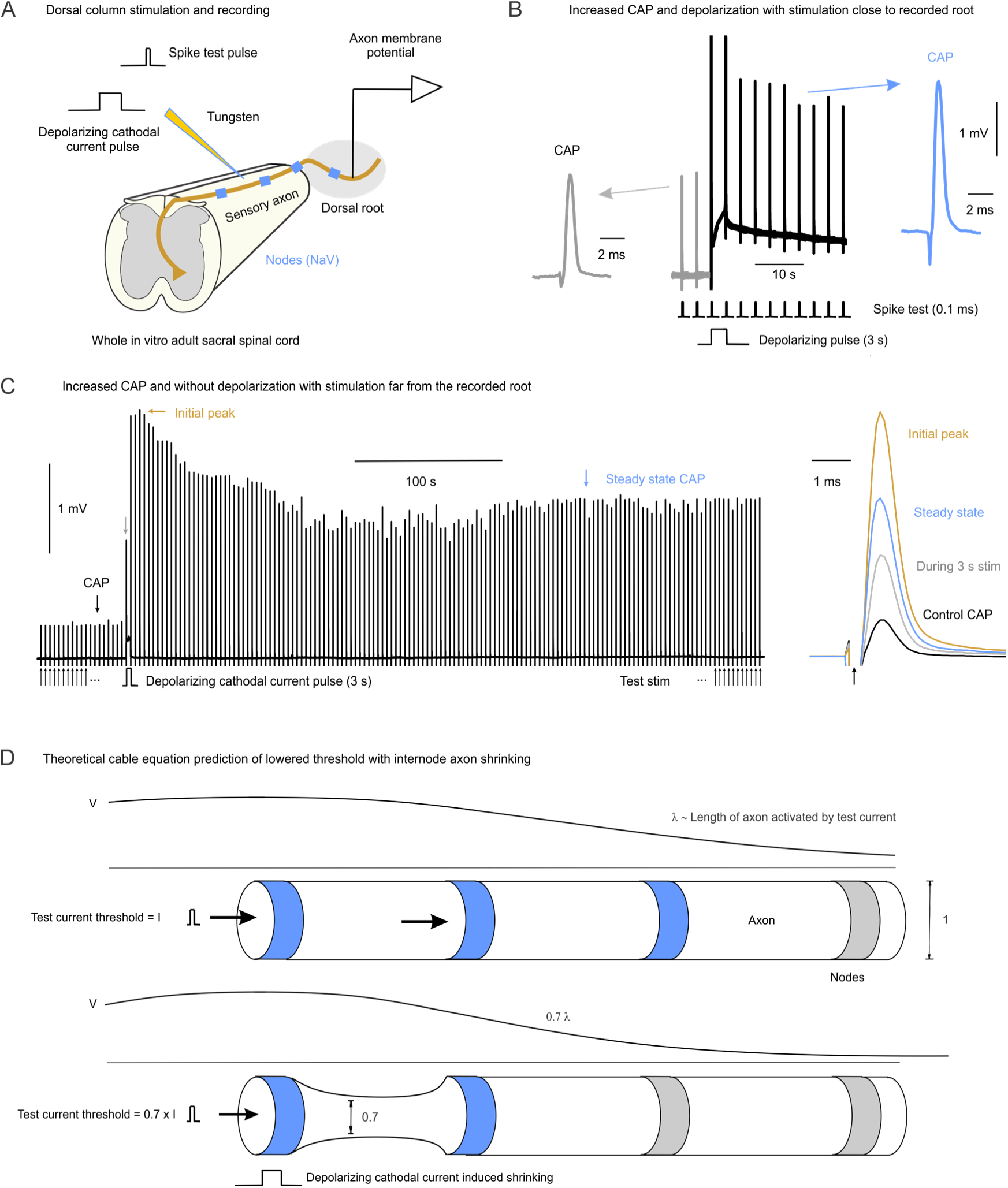
Stimulation raises axon excitability, which cable theory attributes to internode thinning. A, Arrangement for stimulating the dorsal columns of the whole in vitro adult sacral spinal cord with a tungsten electrode, while recording the axon membrane potential and the compound action potential (CAP) evoked by a brief spike test pulse (0.1 ms, 30 μA). B, With the stimulation close to the recorded root, a 3 s depolarizing cathodal pulse (15 μA) increased the CAP evoked by the 0.1 ms spike test pulse and produced a sustained depolarization, with the CAP shown before and after at right. C, With the stimulation further from the recorded root, the same 3 s pulse increased the CAP without a depolarization, with an initial peak that settled to a steady state, and the superimposed CAPs shown at right. Similar results obtained for n = 5 of 5 mice tested in B and C. D, Schematic of cable equation prediction of the membrane potential at the test current threshold needed to evoke a spike. When the internode diameter shrinks by 30% (to 70% of control value) the length constant λ also shrinks by 30%, so that the test current activates a shorter length of axon and the threshold falls, as detailed in the Methods.

Like the reduced axon diameter, the raised excitability remained remarkably constant for long after the stimulation, suggesting that the increased excitability reflects the altered anatomy rather than stimulus-evoked active membrane currents, which often decay over seconds to minutes (as detailed in the next section). Indeed, applying the length-scaling properties of the axon cable equations, described by Stein^2^, to predict the change in threshold current from the anatomical plasticity, the observed increased excitability was accounted for by axon shrinking: the thinned internode presents a higher axial resistance and a shorter length constant, so that the same node is activated by less current, with proportionally less current lost into the adjoining internode (Fig 2D). This can be seen quantitatively from the cable equation for the axon membrane potential V_m_, in which the node carries the membrane current and the internode acts as a passive axial resistor^15^. With the node radius r_n_ fixed and only the internode radius r_i_ shrinking, this axon cable equation is approximated as

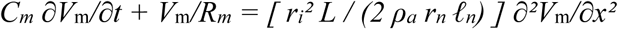

where L is the internode length, C_m_ the membrane capacitance, R_m_ the membrane resistance, ρ_a_ the axoplasmic resistivity, and ℓ_n_ the node length, all fixed constants (see equation (16) in the Methods). Rescaling distance by the internode radius, X = x/r_i_, gives an equation independent of this radius r_i_

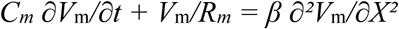

where β is a fixed constant. The voltage profile over distance is therefore universal in the rescaled coordinate, and all of the geometric effect of thinning enters through the stretch x = r_i_ X. Two simple consequences follow. The electrotonic length over which voltage spreads scales linearly with the internode radius, so the length constant falls in proportion to the thinning, and the current needed to bring a node to threshold, set by the fixed node circumference times this spread length^2^, likewise scales linearly with the internode radius. Thus, for the measured thinning to about 70% of the initial diameter, both the length constant and the threshold current fall to about 70% of their initial values, a purely geometric increase in excitability with the node and its channels unchanged. The full derivation is given in Methods. Because the stimulus used to test the CAP was applied near threshold (1.25 to 1.5 times threshold), low on the recruitment curve, this fall in threshold predicts the CAP to increase to 200 to 300% of control (2 to 3 fold increase; see equation (20) in Methods), consistent with the large increases observed here and in Jankowska et al.^6^

In large myelinated sensory axons the electrotonic spread given by the length constant is about twice the internodal distance^16^, as depicted in Fig 2D, which generally prevents failure of conduction between nodes. However, with a reduction of the length constant to 70% of its original value after stimulation we wondered whether this might induce some failure. With our standard 15 to 20 μA stimulation (3 s) this was not an issue with the CAP increasing, rather than decreasing, consistent with a modest reduction in length constant. However, with stimulation closer to the spinal cord (< 50 μm) or larger (25 μA) the CAP was sometimes reduced (n = 7 of 22 trials), though we did not pursue this issue, and left it to future studies related to our earlier investigations of branch point failure^16^.

### A grease-gap signal reports the axial-conductance change of axon thinning

To independently quantify axon shrinking we measured the change in axial resistance with thinning, by inducing a steady current along the axon and measuring the resulting voltage change as this resistance changed. For this, we recorded the composite axonal transmembrane potential by grease-gap methods from the proximal end of a short cut root mounted on a wire in grease close to the stimulation site (Fig 3A), where the recorded voltage V is proportional to the transmembrane potential V_m_ of the axons in the root V = k V_m_, and k is a constant determined only by the axon seal in the grease (k ≈ 0.5, detailed in the Methods). In this arrangement, the spinal cord acts as a current sink and the depolarized cut end as a source, inducing a standing axial current through the internode resistance R_a_, a sealed-end cable boundary effect^17^ detailed in the Methods. The resulting voltage drop from this standing current leaves the axons in the root depolarized relative to the axons deep in the spinal cord that rest at −75 mV^16^, in direct proportion to R_a_ (by Ohm’s law). Thus, a stimulation of the axon in the spinal cord that causes an axon thinning and related increase in the axial resistance should produce a depolarization of the root potential V. Indeed, we observed a significant depolarization with stimulation that we denote dV (0.63 ± 0.43 mV, n = 63 mice, p < 0.05; Fig 3B). This dV developed over the same time course as the axon thinning (steady state peak times of each not different, 10.49 ± 2.65 s, n = 63, p > 0.05), and both were very long lasting (> 1 h), consistent with a common plasticity. Both often did not reverse over the experiment, persisting for hours, with little further thinning or depolarization after a repeated 3 s stimulus (Fig 3B, G). This persistence and lack of within-trial repeatability made the phenomenon difficult to examine before and after drug treatments, without comparing the average dV across differing independent stimulation sites (Fig 3F). Shorter 1 s stimuli, however, produced a partially reversible depolarization about a third as large (×0.31 ± 0.12, n = 7) and lasting only a minute, which we used to estimate the full dV expected at a given dorsal column site with our standard 3 s stimulus, enabling us to predict the dV expected when a drug was applied between the 1 and 3 s stimuli (Fig 3B, C; by multiplying the 1 s stimulation by 1/0.31 ≈ 3). This and inter-trial comparisons allowed for evaluating dV changes with drug manipulations, detailed below (Supplementary Table 1).

**Figure 3.**
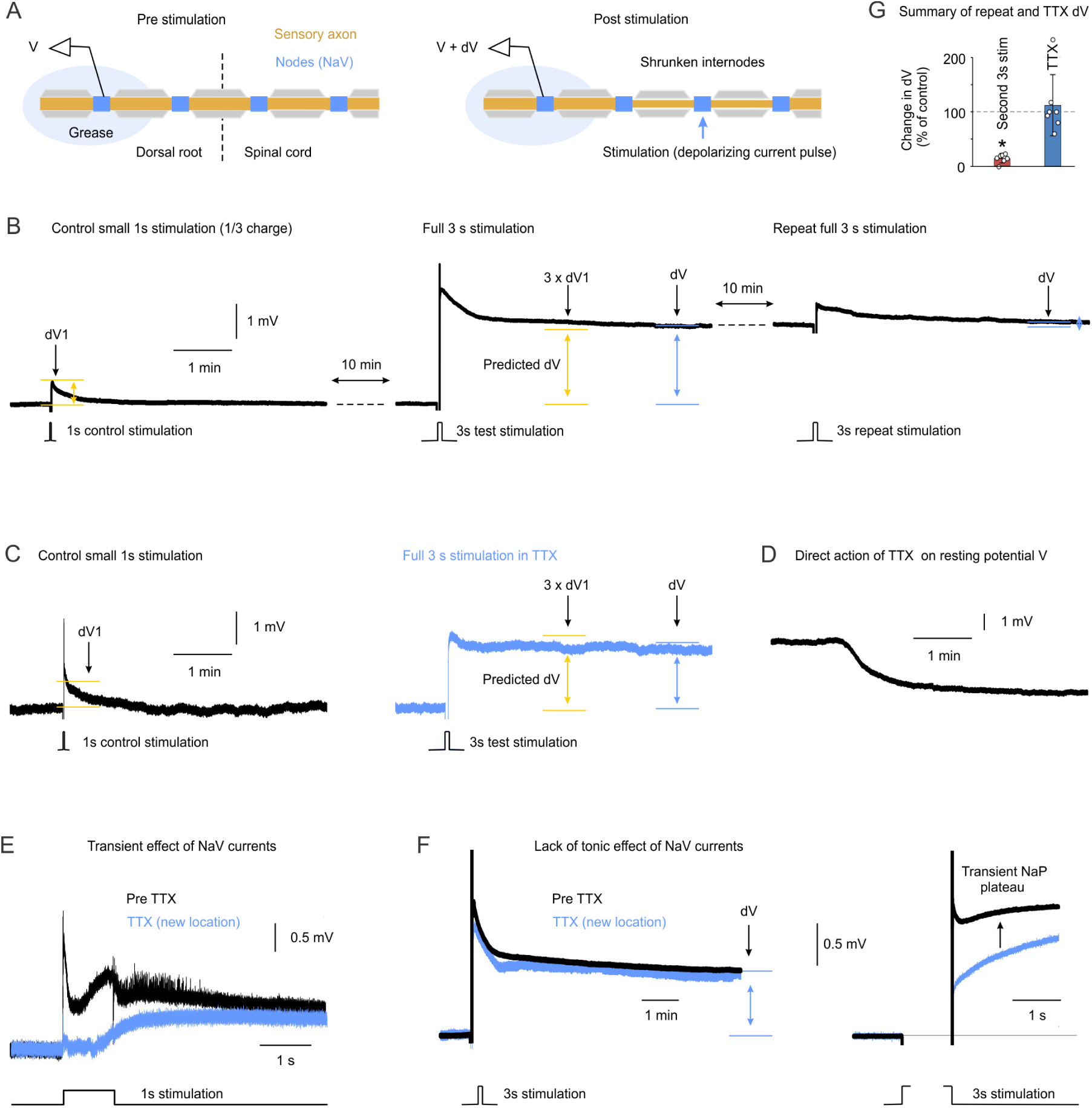
Grease-gap recording of the shrinkage-related dV. A, Recording arrangement, with the dorsal root mounted in grease and the recorded potential V rising to V + dV as the internodes shrink after stimulation in the spinal cord. B, A small 1 s control stimulation (20 μA, < T = 25 μA, cathodal) evokes a transient depolarization of amplitude dV1, whereas a 3 s stimulation (20 μA) evokes a sustained dV with amplitude well predicted by 3 × dV1. A repeat 3 s stimulation 10 min later evokes little further dV. C, Standard drug testing protocol. The same protocol as B but applying TTX (2 μM) after the small 1 s control stimulation, showing that the sustained dV persists without voltage-gated Na currents. D, Direct action of TTX on the resting potential V. E, Transient effects of NaV currents during a 1 s stimulation, before and after TTX. F, Alternative protocol also showing lack of a tonic effect of NaV currents on the sustained dV, with the transient persistent Na plateau shown at right, but in this case the responses are from stimulation of two independent sets of axons, one before and the other after, the former using the sequence in B to assure a comparable setup. G, Group changes in dV with repeated 3 s stimulation of B (n = 7) and with TTX application (n = 7), the latter using both protocols of C and F. * significant change relative to control (100%), p < 0.05.

Axons exhibited persistent transmembrane currents induced by the stimulation that contributed a depolarization, but these generally decayed over the first few seconds after stimulation, and thus were unrelated to the lasting structural change. For example, a sodium current (NaP) evoked by stimulation produced a transient overshoot in dV that was blocked by TTX (Fig 3E, F) while the sustained, shrinkage-related dV was unchanged (Fig 3C-F). To mostly avoid these currents, we only quantified the dV at 10 min after stimulation, well into the steady state anatomical change. The grease-gap dV thus offers a simple assay of axon thinning without the need to image axons in every experiment. To confirm this we exhaustively tested many possible persistent currents that might contaminate dV and failed to find any, as detailed in subsequent sections, and thus we conclude that the sustained dV reflects predominantly the axon diameter reduction.

The dV grows steeply when axons shrink because it is proportional to the change in axial resistance, which is inversely proportional to the axon cross-sectional area or squared radius. Thus, shrinking the radius by a factor f raises the resistance to 1/f² of its resting value, so the change in resistance, and thus the dV, grows in proportion to (1/f² − 1). Quantitatively, this predicts that under reasonable assumptions, a depolarization of about 10 mV occurs in a stimulated axon for the observed thinning to 70% of the diameter (Methods, equation (13), where f = 0.7). Only a fraction (∼10%) of the axons in a root recorded by the grease gap were stimulated, the remaining diluting the signal, and the leak in the grease gap further reduced the recorded signal (k = 0.5). Thus, after correcting the recorded dV for these factors it was close to that predicted (12.67 ± 8.62 mV, 20× the recorded value), though the exact correction does not affect our use of dV as a readout of axon plasticity.

Stimuli that evoked a sustained dV were up to an order of magnitude lower than the spike threshold (T, which was typically about 25 μA), as long as the product of the current and the stimulus duration exceeded a critical value of about 60 μA s (60 s at 1 μA, 15 s at 4 μA, and 3 s at 20 μA all produced a dV, n = 5 of 5 tested each), similar to the reported currents required to increase axon excitability^6^. This implies that the total charge or ion movement is the critical factor driving the plasticity, rather than simply the activation of a voltage-gated current, suggesting a buildup of ions beyond a critical threshold and consistent with an osmotic mechanism, as suggested by Trigo and Smith^4^ and elaborated below.

### GABA_A_ receptor activation drives the same axon thinning

Because GABA, glycine and other transmitter receptors are expressed on sensory axons and directly activate nodes in and around the dorsal columns^16,18,19^, we asked whether the same structural plasticity seen with electrical stimulation is triggered physiologically by GABA and glycine. Transient application of GABA and glycine or the GABA_A_ receptor agonist muscimol, for 7 min, produced long-lasting axon thinning, maintained for the duration of the recording (> 1 h; to about 72% of control; Fig 4A, B), perhaps contributing to the increased excitability and tonic primary afferent depolarization that outlasts GABA_A_ receptor activation in these axons^16^. This GABAergic and glycinergic activation also produced a long-lasting depolarization that developed with the long-lasting axon thinning, which we termed dV_g_ (Fig 4C; 0.33 ± 0.22 mV for GABA alone, 1.03 ± 0.23 mV for muscimol, and 0.73 ± 0.20 mV for GABA + glycine together, 7 min each, with n = 10, 16 and 16 mice respectively, significant depolarizations, p < 0.05), consistent with shrinking and an increased axial resistance. Optogenetic activation of GABAergic neurons that innervate sensory axons for 1.5 min (in GAD2//ChR2 mice, 12 Hz train of 10 ms light pulses, 0.7 mW/mm^2^) produced a similarly long lasting dV_g_ (0.49 ± 0.08 mV, n = 5, p < 0.05). Critically, the sustained dV_g_ and axon thinning occluded the response to direct nodal stimulation dV (Fig 4C; Supplementary Table 1K; Supplementary Fig 1 bottom centre), indicating that the two share the same mechanism and the same structural endpoint. This action of GABA likewise lasted hours, with the ability of a stimulus to evoke a dV recovering only after about six hours (Fig 4C; Supplementary Table 1K, last rows). However, dV_g_ itself decayed slowly over an hour, suggesting that GABA generated more moderate plasticity somewhere between the dV of a 1 and 3 s cathodal stimuli (Fig 3B).

**Figure 4.**
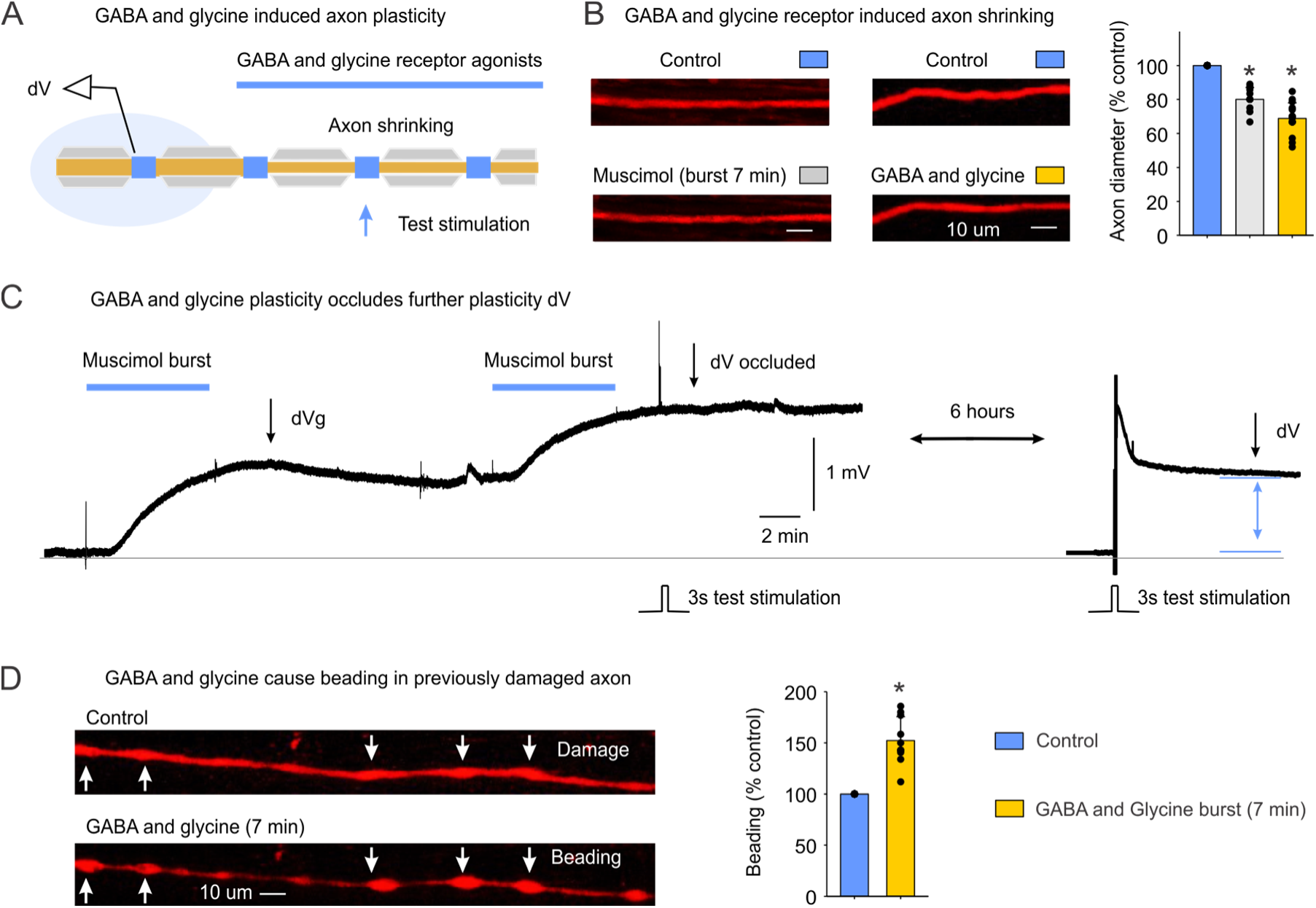
GABA and glycine receptor activation produces the same axon plasticity that mimics stimulation. A, Arrangement for bath application of GABA and glycine receptor agonists while recording the axons, with the test stimulation applied to the dorsal columns (20 μA, 3 s). B, Live imaging of axons before and after the GABA_A_ receptor agonist muscimol (15 μM, 7 min burst) or GABA and glycine together (2 mM each), with group data showing the reduced axon diameter (percent of control). Axons labelled with tdTom (red) rather than GFP in this case. * significant change, p < 0.05, n = 10 and 15 respectively. C, A muscimol burst produced a sustained depolarization dV_g_ and occluded the dV evoked by a subsequent 3 s test stimulation, with the dV recovering only after about six hours (right). Similar results in n = 5 of 5 mice. D, In previously damaged axons, GABA and glycine produced beading (arrows), with group data showing the increased bead diameter (percent of control). * significant change, p < 0.05, n = 10.

### Motor axons show the same activity-dependent axon thinning

We next enquired whether the anatomical plasticity seen in sensory axons also occurred more generally in other central axon types. For this we stimulated motor axons at the ventral root entry zone of the spinal cord with the same subthreshold cathodal pulse (tungsten electrode, 3 s, 20 μA) and recorded the shrinkage-related dV on the ventral roots. As with sensory axons, we observed a prominent long lasting dV on motor axons (0.84 ± 0.51 mV, n = 5 mice, significant change, p < 0.05), suggesting that these axons exhibit similar anatomical plasticity to sensory axons.

### Axon thinning is initiated by under-myelin Kv1 currents, the Na/K-ATPase and glial K⁺ clearance

Having developed an assay of axon thinning dV, we used it to dissect the underlying mechanisms of axon plasticity with pharmacological blockers, which is otherwise difficult to do by excitability and imaging alone. To start, we screened many compounds to determine which channels, receptors or transporters were involved in initiating the shrinkage-related dV, with all results summarized in Supplementary Table 1, and associated Supplementary Fig 1.

From the outset we noticed that the sustained dV was unaffected by blocking voltage-gated Na channels underlying the spike with TTX (Fig 3C-F), even at very high doses that block TTX resistant NaV (Supplementary Table 1A). Likewise, dV initiation was unaffected by blocking fast synaptic transmission involving glutamate, GABA or glycine receptors with CNQX, APV, gabazine and strychnine (Fig 5B, G; Supplementary Table 1H). This enabled us to study other channels underlying the origins of the plasticity in axons isolated by these blockers from synaptic inputs or the many currents evoked during spiking, as in the following sections.

**Figure 5.**
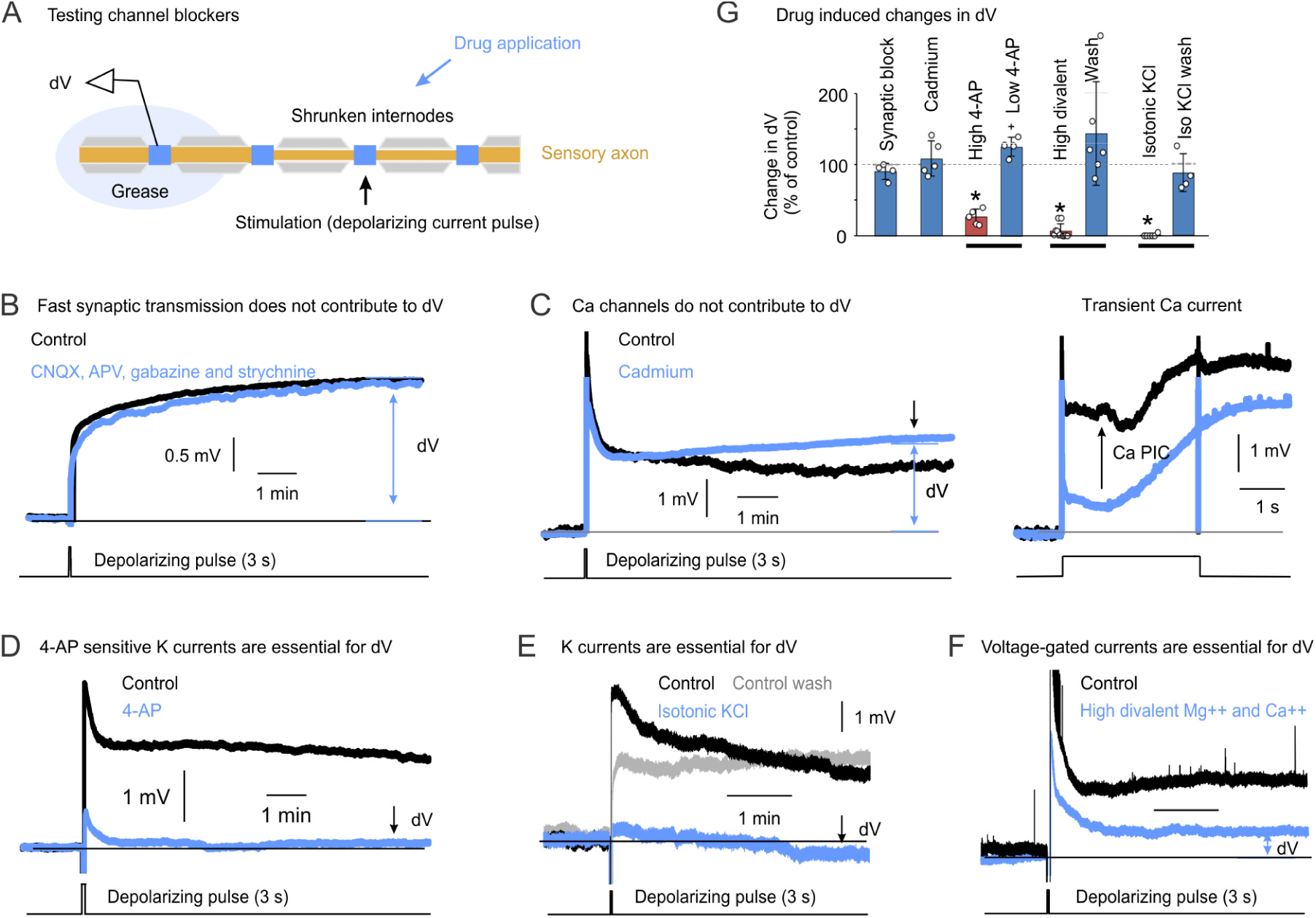
Voltage-gated potassium currents are necessary for the axon thinning dV. A, Recording arrangement, as in Figure 3, with the dorsal root in grease and the recorded potential rising to V + dV as the internodes thinned after stimulation of the spinal cord. B, Blocking fast synaptic transmission with CNQX, APV, gabazine and strychnine did not change the dV evoked by a 3 s subthreshold stimulation (20 μA), using the protocol of Figure 3F. C, Blocking calcium currents with cadmium (400 μM) did not change the dV, although it removed a calcium current early in the response. The right panel shows this on an expanded time scale, where persistent calcium currents (Ca PIC) were triggered by the stimulation but decayed within seconds. D, Blocking Kv1 currents with a high dose of 4-AP (1 to 3 mM over more than 1 h) abolished the dV. E, Replacing the ACSF with isotonic KCl abolished the dV, which returned on washing back to normal ACSF. F, Inhibiting voltage-gated currents with high divalent cations (5 mM Ca and 15 mM Mg added, with equimolar Na removed to maintain osmolarity) inhibited the dV. G, Summary bar graphs of the drug induced changes in dV in B to F, expressed as a percentage of control. * significant decrease and + significant increase, p < 0.05. n values are given in Supplementary Table 1. C to F were recorded in TTX (2 μM), CNQX (50 μM), APV (50 μM), gabazine (50 μM) and strychnine (5 μM).

Blocking voltage-gated Ca²⁺ channels did not prevent initiation of the dV (Fig 5C, G; Supplementary Table 1B cadmium), nor did removing bath Ca²⁺, chelating Ca²⁺ with EGTA, blocking ryanodine receptors on the endoplasmic reticulum or blocking mPTP on mitochondria (Supplementary Table 1B). Blocking the persistent inward cation current I_h_ also did not reduce dV (Supplementary Table 1A). Blocking TRP related non-selective cation currents also did not reduce dV (Supplementary Table 1A). Thus, while we had initially thought a depolarizing voltage-gated channel might be essential to initiating dV, none were. Blockers of connexins, ACh receptors, purinergic receptors, PKA, astrocyte function, and most transporters or exchangers (Supplementary Table 1F, H, I, and J) were also without effect in blocking the initiation of dV.

We thus turned to K currents which seemed counterintuitive as these are inhibitory. Indeed, blockers of the K currents mediated by K2P, Kv7 and Kir were without effect (Supplementary Table 1C). However, blocking voltage-gated K⁺ channels with 4-aminopyridine (4-AP) prevented the stimulus-induced dV (Fig 5D, G). The doses required were high (Supplementary Table 1C), consistent with an action on Kv1 channels beneath the myelin, where comparable doses are needed to reach these channels during direct periaxonal recording^20–22^. Collapsing the K⁺ gradient with isotonic KCl likewise blocked a stimulus from inducing a dV, and a second stimulus delivered after washout of KCl again evoked a dV, showing that stimulation alone, without K⁺ currents, does not induce the structural change (Fig 5E, G; Supplementary Table 1C). Inhibiting voltage-gated channels by increasing surface-charge screening with high divalent cation concentrations^23^ also inhibited the stimulus-induced dV, and again a divalent washout enabled a second stimulation to evoke a full dV (Fig 5F, G; Supplementary Table 1C). Low doses of 4-AP tended to slightly increase dV, in contrast to high doses (Fig 5G; Supplementary Table 1C), consistent with Kv1 channels near the node being reached first and opposing the trigger for axon thinning from Kv1 under the myelin. Thus, overall a voltage-gated K current carried by 4-AP sensitive Kv1 channels under the myelin is involved in initiating the axon plasticity.

These results point to a build-up of K⁺ beneath the myelin as the trigger for axon thinning, as detailed in the Fig 6A schematic. Barrett and colleagues showed that K⁺ released from Kv1 channels accumulates in the bounded periaxonal space and can drive regenerative K⁺ currents^20,21,24^. Such accumulation, with its accompanying osmotic load, may compress the axon and swell the surrounding glia. The osmotic load is amplified by the Na/K-ATPase pump, which is distributed along the internode with Kv1 ^22,25,26^ and moves three Na⁺ into the periaxonal space for every two K⁺ it retrieves^23^. The pump is also required to maintain the steep glial K⁺ gradient that lets inwardly rectifying Kir4.1 channels siphon K⁺ from the periaxonal space and swell the glia^27–29^, with connexin (Cx29) hemichannels in the innermost myelin providing a further path from the periaxonal space into the oligodendrocyte^22^. Blocking the pump with ouabain abolished the shrinkage-related dV (Fig 6B, E; Supplementary Table 1E), implicating the Na/K-ATPase in this structural plasticity, as Trigo and Smith^4^ concluded for high-frequency firing. The block by ouabain required relatively lower doses than the block of under-myelin Kv1 by 4-AP, suggesting that some of the ouabain effect is on the bath-accessible Na/K-ATPase of the glia, which supports the Na⁺-dependent water transport and swelling that accompany uptake of the periaxonal K⁺ load.

**Figure 6.**
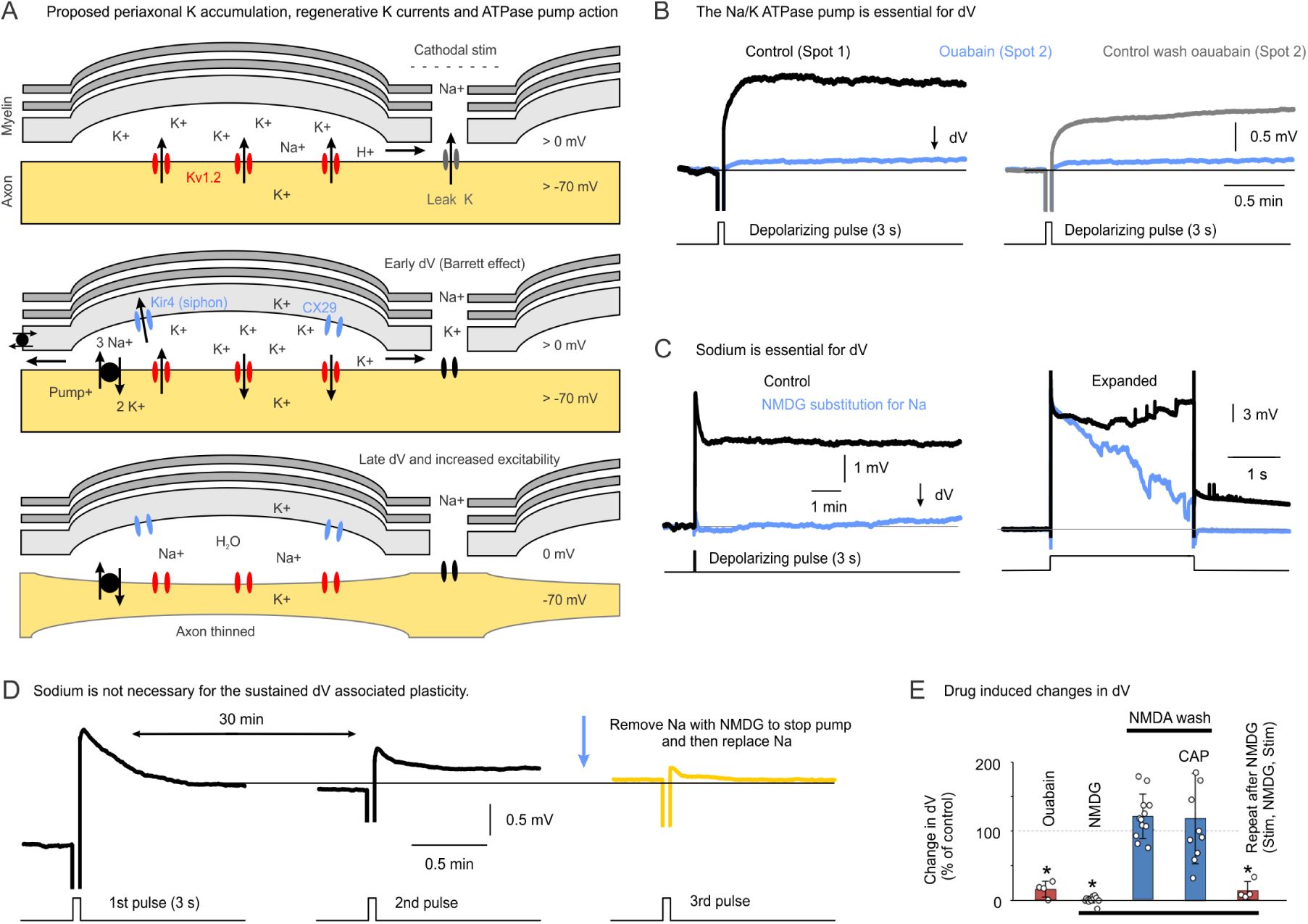
The Na/K-ATPase pump and sodium are required to initiate the axon thinning. A, Proposed model of axon thinning with cathodal stimulation. Kv1 currents open beneath the myelin and release K, which accumulates in the periaxonal space and regeneratively drives further Kv1 opening (Barrett effect), while the Na/K-ATPase pump amplifies the periaxonal ion build-up, and glial Kir4.1 siphoning and connexin (CX29) coupling carry K⁺ into the glia and swell them, culminating in osmotic thinning of the axon. B, The dV evoked by stimulation (20 μA, 3 s at control spot 1) was blocked by inhibiting the pump with ouabain (100 μM, spot 2), and returned on washout (spot 2, wash). C, The dV was likewise blocked by removing the pump substrate Na and replacing it with NMDG. NMDG also blocked a transient persistent sodium plateau (NaP), shown on an expanded time scale at right to show onset of dV. D, Once the axon thinning dV had been evoked (first pulse), it was not reversed by transiently blocking the pump with NMDG and then restoring Na (third pulse), showing that the pump and sodium are needed to establish the thinning but not to sustain it. E, Summary bar graphs of the changes in dV with ouabain (B), with NMDG substitution for Na (C), after washout of NMDG (change in CAP also shown), and with a repeat stimulation before and after NMDG (D), expressed as a percentage of control. * significant decrease, p < 0.05. n values are given in Supplementary Table 1. B, C and D were recorded in TTX (2 μM), CNQX (50 μM), APV (50 μM), gabazine (50 μM) and strychnine (5 μM).

Indirectly blocking the Na/K-ATPase pump by removing its substrate Na⁺ from the bath by replacing Na⁺ with NMDG⁺ also blocked the dV (Fig 6C, E; Supplementary Table 1E). This NMDG⁺ action was faster and more thorough than with ouabain, suggesting that additional Na/K-ATPase were blocked, likely under the myelin where small ions like Na⁺ and NMDG⁺ more readily move. Also, a stimulus that failed to evoke a dV in NMDG⁺ or ouabain induced a dV (or increased CAP) after NMDG⁺ or ouabain were washed out, confirming that Na⁺ and the pump are needed for axon thinning and that the stimulus does not otherwise produce plasticity (Fig 6B, E; Supplementary Table 1E). Replacing Na⁺ with NMDG⁺ after the dV had been induced did not reverse the thinning, since a repeat stimulus at the same spot after restoring Na⁺ did not evoke a dV of similar size (Fig 6D, E; Supplementary Table 1L), showing that the pump and Na⁺ are needed to establish the thinning but not sustain it.

We next examined whether a thinning-related axial resistance change associated with dV could be triggered by depolarization of just one axon, by employing direct intracellular current injection into single axons. Large 30 mV depolarizations lasting 3 to 10 s to well over spike threshold produced no sustained change in resistance or membrane potential (1.1 ± 5.2% or 2.3 ± 8.3%, respectively, n = 5 each, p > 0.05). Thus, our standard 3 s extracellular depolarization of axons that caused axon thinning (Figs 1-5) likely relied on simultaneous activation of many nearby axons that overwhelmed the extracellular K^+^ handling.

### A coincident local alkalinization helps subthreshold stimuli or GABA engage Kv1

We next inquired how a subthreshold depolarization so effectively activates K^+^ current-mediated axon thinning when Kv1 channels are normally only partially activated at these potentials. One possibility is that a pH shift from electrochemistry at the tungsten electrode (cathode) contributes, perhaps lowering K^+^ channel activation thresholds. The cathode can produce OH^−^ by numerous Faradaic reactions, including oxygen reduction (O_2_ + 2H_2_O + 4e^−^ → 4OH^−^) or electrolysis of water (2H_2_O + 2e^−^ → H_2_ + 2OH^−^), with the latter H_2_ likely able to disperse before forming bubbles at the low currents we used^30^. Employing a pH-sensitive microelectrode, we found that our usual 3 s subthreshold stimulus produced a transient local alkaline shift from pH 7.4 to on average about pH 7.9 over 10 s at our standard axon distance of 50 μm from the electrode. This pH shift decreased progressively over a 200 μm distance from the electrode (Fig 7).

**Figure 7.**
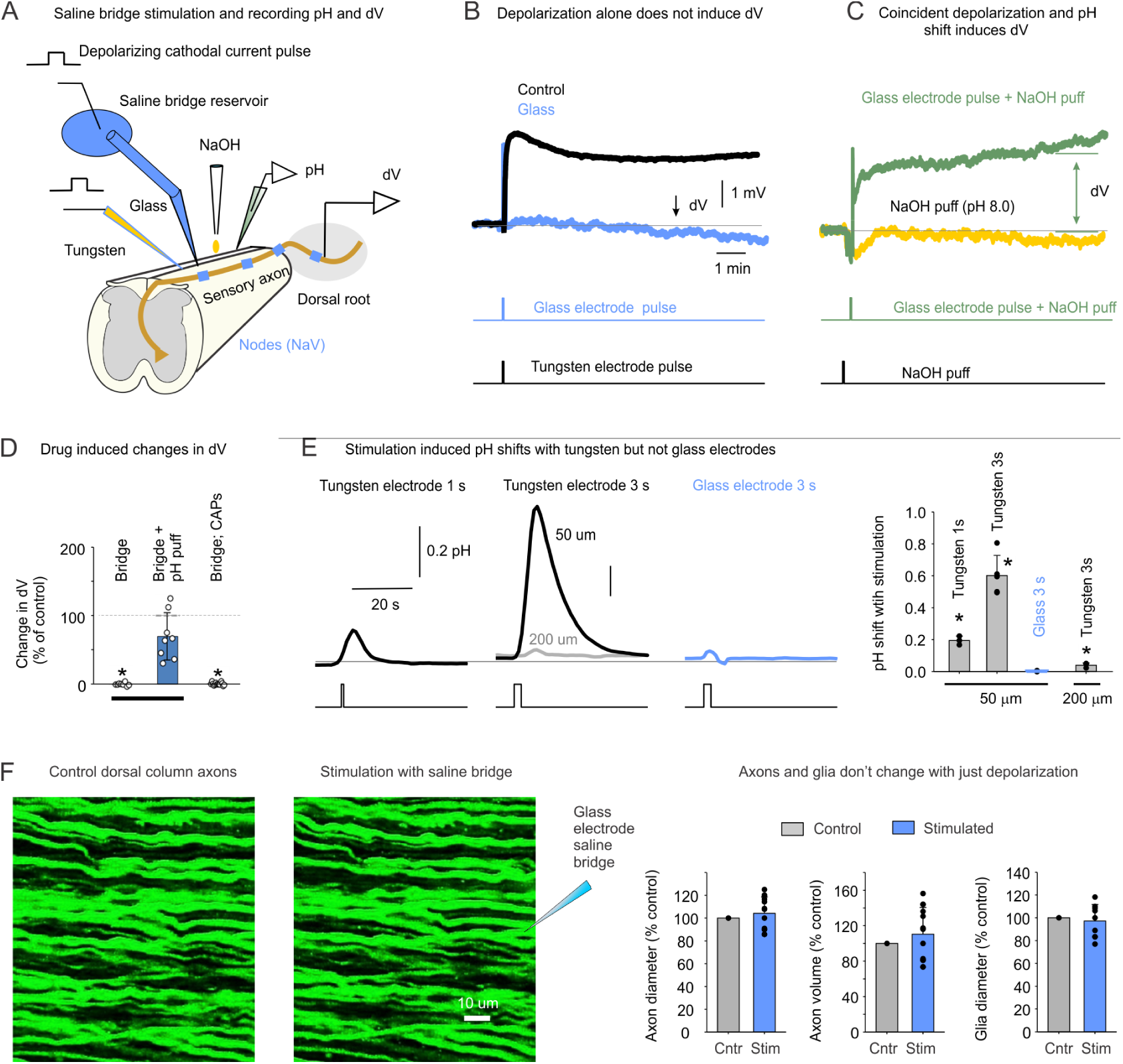
Axon plasticity requires a coincident depolarization and alkaline shift. A, Arrangement for stimulating the dorsal columns through a saline bridge with a glass electrode, which depolarizes the axons without the electrochemistry of a metal electrode, or with a tungsten electrode, while recording the local pH and the root potential. B, Depolarization alone did not induce a dV. A tungsten electrode pulse (20 μA, 3 s) evoked the usual dV (control, black), whereas the same depolarization delivered through the glass electrode and saline bridge did not (blue). C, Coincident depolarization and alkalinization induced a dV. A NaOH puff alone (pH 8.0) did not evoke a dV (yellow), whereas the glass electrode pulse combined with the NaOH puff did (green). D, Summary bar graphs of the changes in dV and CAP with saline bridge stimulation alone, and of the dV rescued by combining bridge stimulation with a NaOH puff to pH 8.0, expressed as a percentage of control. * significant decrease, p < 0.05. n values are given in Supplementary Table 1. E, Stimulation induced local pH shifts with tungsten but not glass electrodes, for both 1 s and 3 s pulses (20 μA). The pH was recorded 50 μm from the electrode in mACSF in all conditions, except when 200 μm indicated. Group data at right show the pH shift with stimulation for each condition. * significant change, p < 0.05. F, Live imaging of dorsal column axons before and after stimulation through the glass electrode and saline bridge, with group data showing that the axon diameter, axon volume and glia diameter were unchanged (percent of control), confirming that depolarization alone does not produce the anatomical plasticity.

To separate the stimulus-induced depolarization from the electrochemistry, we stimulated through a saline bridge, where the electrode was placed in a large HEPES-buffered reservoir that diluted any pH shift and this was connected to a micropipette that delivered current to the axons (Fig 7A). Applying our standard 3 s depolarization stimulation through this saline bridge increased the CAP during the stimulation (296.1 ± 159.4% increase, n = 5 mice), demonstrating that the bridge can readily depolarize the axons. However, this bridge stimulation did not produce a sustained dV or change in the axon diameter or CAP (Fig 7B, D, F; Supplementary Table 1G), showing that depolarization alone is not sufficient for the plasticity. Raising the in vitro bath pH to 8.0 with transient NaOH (puff) was likewise insufficient alone to produce a sustained change in potential V (Fig 7C; −0.25 ± 0.32 mV, n = 6, p > 0.05). Combining the saline-bridge depolarization with the pH shift, however, did induce a shrinkage-related dV (Fig 7C, D; Supplementary Table 1), indicating that coincident depolarization and alkalinization are required. The alkaline shift may act through modulating the Kv channel sensitivity or a related channel like K2P, though we did not pursue this further. A similar saline bridge stimulation applied in vivo also did not increase the CAP (data kindly provided by E. Jankowska and I. Hammar, n = 1).

Notably, GABA_A_ receptor activation also alkalinizes the extracellular space as it depolarizes the axon through bicarbonate (HCO_3_⁻) as well as Cl⁻ efflux^31–34^, suggesting that a shared local alkalinization may accompany both subthreshold stimulation and the physiological action of GABA on axon thinning. Reducing bicarbonate with acetazolamide and replacing standard bicarbonate-buffered ACSF with HEPES buffered ACSF partially reduced the stimulus-evoked dV and completely blocked the GABA-evoked dV_g_ (Supplementary Table 1G and Supplementary Fig 1), consistent with bicarbonate movement sustaining the local pH shift that triggers axon thinning, with the GABA route depending most heavily on bicarbonate efflux.

### Clinical high frequency stimulation protocols produce the same axon thinning

While we have to this point deliberately employed weak cathodal pulses to induce beneficial axon plasticity similar to that with GABAergic activation, we next examined whether stimulation protocols used clinically might act similarly. For this we started by testing suprathreshold high frequency stimulation trains of short pulses over 1 min (100 Hz, 30 μA, 0.2 to 0.4 ms pulses, 1.5×T), though we did this in the presence of TTX to focus on local node effects (T measured before TTX). This too produced a sustained dV (0.53 ± 0.49 mV, n = 6 mice, p < 0.05; Fig 8C, left), indicating axon thinning. Charge-balanced pulse trains that more closely mimic clinical applications likewise produced axon thinning, though a comparable dV required more total cathodal charge, with longer trains and larger pulses (∼3 min, 100 Hz, 100 μA, 1 ms cathodal pulse followed by 1 ms anodal; 0.46 ± 0.29 mV, n = 7, p < 0.05; Fig 8C), and slight errors in the charge balance induce larger dV values (not shown). Popular 10 kHz charge-balanced stimulation protocols^35^ readily produced a shrinkage-related dV, because of the very large cathodal duty cycle (33% and charge movement), fast switching, and the slow irreversible nature of the Faradaic reactions involved (1.5 min, 10 kHz, 30 μA, 0.033 ms cathodal followed by 0.033 ms anodal; 0.64 ± 0.55 mV, n = 6, p < 0.05; Fig 8C). These trains were again related to a pH shift because they produced an ∼0.5 pH shift at the tip of the stimulating electrode (Fig 8B). An alkaline pH shift was observed over the range of pulse widths commonly used for stimulation trains (Fig 8D), and also occurred when the tungsten electrode was replaced with the 30 μm fine Pt-Ir wires used for spinal cord stimulation (Fig 8E)^36–38^. Thus, high frequency stimulation over a long period can induce similar plasticity to physiological GABA_A_ receptor activation, and might contribute to clinical effects of spinal cord stimulation.

**Figure 8.**
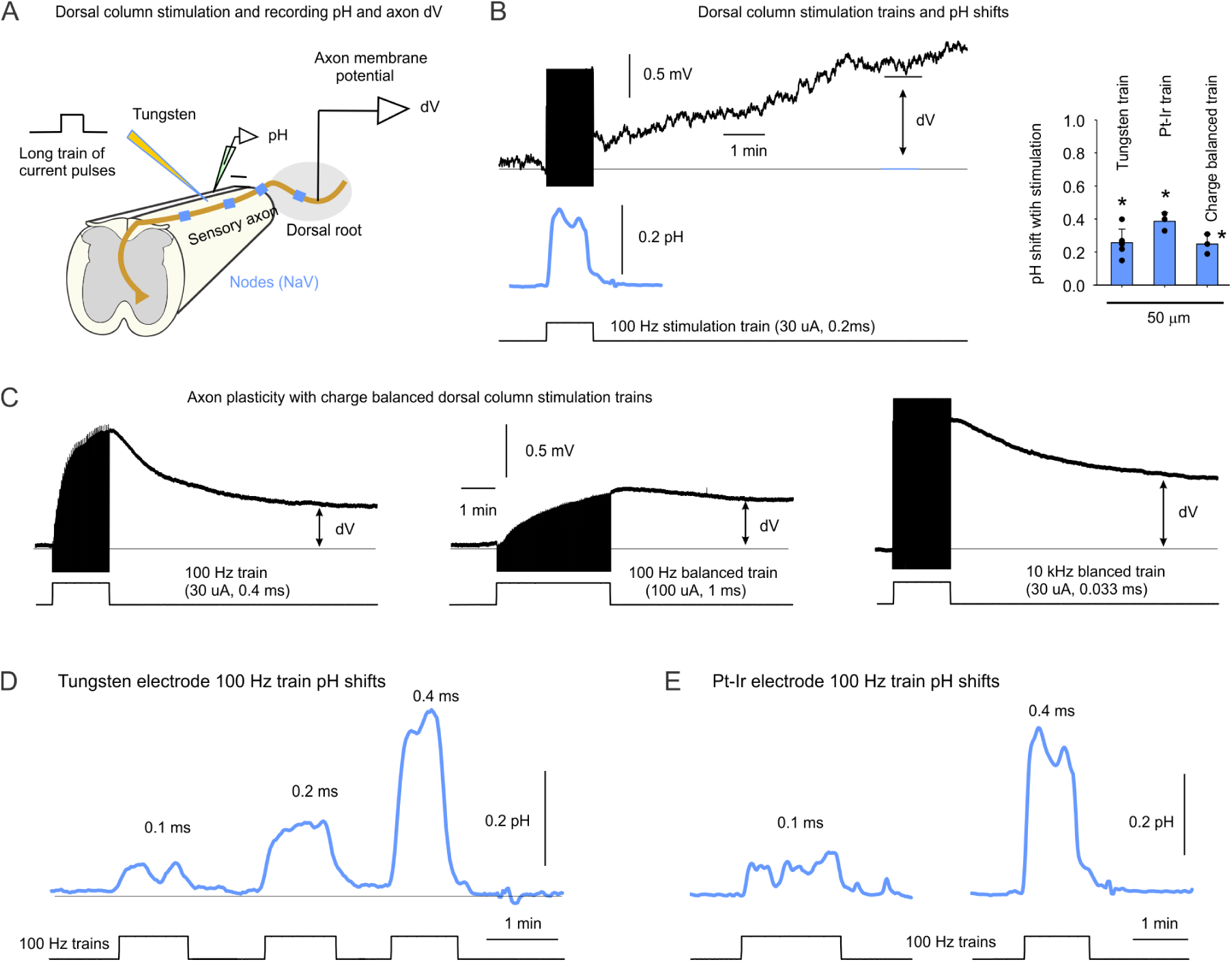
High frequency stimulation trains produce a local alkalinization and a sustained dV. A, Arrangement for applying a suprathreshold depolarizing current pulse train to the dorsal columns with a tungsten electrode, while recording the local pH with a pH microelectrode and the axon membrane potential from the dorsal root. B, A 1 min train of 30 μA 0.2 ms pulses at 100 Hz produced a sustained depolarization dV, indicating axon plasticity, with the simultaneously recorded local pH shift below. Group data at right show the pH shift with tungsten, Pt-Ir and charge-balanced trains, recorded 50 μm from the electrode. * significant change, n = 3 to 5 each, p < 0.05. C, Axon plasticity with charge-balanced trains. A 100 Hz train (30 μA cathodal pulse, 0.4 ms), a charge-balanced 100 Hz train (100 μA, 1 ms cathodal followed by 1 ms anodal) and a charge-balanced 10 kHz train (30 μA, 0.033 ms cathodal followed by 0.033 ms anodal) each produced a sustained dV. D, Local pH shifts recorded 50 μm from the tungsten electrode during the 100 Hz trains, which grew with the pulse width (0.1, 0.2 and 0.4 ms). E, Similar pH shifts with a platinum-iridium (Pt-Ir) electrode rather than tungsten, at 0.1 and 0.4 ms pulse widths, indicating that the alkalinization does not depend on the electrode metal. Similar results obtained in n = 5 of 5 tests for D-E.

### Water and anion movement occurs through multiple routes

Cell-volume change from cation movement requires matched movement of water and anions. No single blocker to water-transporters (AQP4, NKCC1 or KCC2; Supplementary Table 1F) or Cl⁻-channels (GABA_A_, GlyR, VRAC and CLC2; Supplementary Table 1D) eliminated the shrinkage-related dV, indicating that water and anions move through several parallel routes, perhaps including the incompletely sealed paranodal junction where the myelin meets the axon^39,40^. Replacing most bath Cl⁻ with gluconate did not block the dV (Supplementary Table 1D; Supplementary Fig 1), suggesting that HCO_3_⁻ can substitute for Cl⁻ in the osmotic movement. The anion-exchanger (AE) blocker DIDS increased the dV (Supplementary Table 1F; Supplementary Fig 1), consistent with AE normally dissipating the bicarbonate and osmotic gradient, through its exchange of bicarbonate with Cl⁻, ^41^ so that blocking it intensifies the shrinkage or spreads the pH shift to more axons in the recording (though DIDS is not selective and also blocks other anion transporters and channels).

### Axon thinning is not reversed once established

Once shrinkage had occurred, no compound reversed the stimulus-induced dV or increased excitability, when applied long after stimulation (> 10 min). That is, when compounds like NMDG and isotonic KCl that block the initiation of shrinking were added after a shrinkage-related dV or change in CAP was evoked, then after removing the compound a subsequent stimulus did not evoke a dV, as mentioned above (Fig 6D, E; Supplementary Table 1). Thus, there was no re-dilation of the axon that rescued its ability to shrink again. Furthermore, most compounds alone caused a direct depolarization rather than hyperpolarization, not reversing the dV (Supplementary Table 1). A few compounds like TTX hyperpolarized axons (Fig 3D; Supplementary Table 1A) but these did not reduce dV. ZD7288, which blocks I_h_ HCN currents, also caused a hyperpolarization of the axons but tripled a subsequent stimulus-evoked dV (Supplementary Table 1A), implying that loss of the stabilizing action of I_h_ leads to more axons shrinking and/or greater shrinking. Such large dV values made us wonder whether this affected spike transmission or damaged the axon. Indeed, we observed a slow decay of the CAP amplitude over an hour after I_h_ was blocked, though we did not image axons in ZD7288 and left this open for future studies.

### Mechanical weakening converts axon thinning into pathological beading

In some preparations a few axons were inadvertently weakened during dissection, by slight stretch of the roots or contact with the electrode, visible as small enlargements along the axon. In these axons, GABA and glycine produced not only local thinning but large-scale swelling at the weakened sites (Fig 4D), culminating in frank beading, the precursor to irreversible damage including axotomy and spheroid formation^42–44^. The physiological process that normally thins axons and raises their excitability can therefore, where axons are mechanically compromised, drive them instead toward damage.

## DISCUSSION

Our results demonstrate that focal subthreshold stimulation or GABAergic activation of central myelinated axons drives a rapid, long-lasting thinning of the stimulated internode, matched by swelling of the surrounding glia, that leaves the node, paranode and more distal axon segments relatively unchanged. The thinned axon is markedly more excitable, and both this and the anatomical plasticity persist for hours. The magnitude of the excitability change is quantitatively accounted for by cable theory: a thinner internode has higher axial resistance and a shorter length constant, so that less stimulating current is lost along the axon and the node reaches threshold more easily^2^. The higher axial resistance also leads to an increased depolarization from currents passed axially, dV. Both the sustained increase in excitability and dV are accounted for predominantly by the altered axon geometry because we found no sustained membrane current that contributes over the hours-long time course of these effects. Overall, this provides a stable, lasting form of axonal plasticity from a brief stimulation of a node or physiological activation of GABA_A_ receptors on a node, which occurs in sensory and motor axons.

The axon thinning is initiated by voltage-gated potassium currents, likely through Kv1 channels beneath the myelin, and requires the Na/K-ATPase pump, bicarbonate, a pH shift, and altered potassium clearance. We interpret this as a regenerative build-up of potassium in the bounded periaxonal space: Kv1 opening releases K⁺ that accumulates^20,21^ and with the osmotic load amplified by the pump, draws water and anions to compress the axon while the glia swell to take up the space through Na⁺-dependent water transport^28^. The low-dose block by ouabain points to a bath-accessible glial pump contributing, since impairing it prevents the glia from swelling to accommodate the potassium load. However, removal of Na (and replacement by NMDG) more thoroughly blocked the shrinkage related dV, consistent with the pump under the myelin also being involved, being more accessible to small ions like this. This mechanism parallels the osmotic, pump-dependent internode thinning that Trigo and Smith^4^ described after high-frequency firing of peripheral axons, and extends it to subthreshold activation of central axons, where the structural change raises rather than impairs excitability and spares the node. Related activity-dependent remodelling of the node and paranode has been imaged directly by others, including paranodal myelin retraction with high-frequency stimulation^3^ and swelling of the node and paranode^4,5^. Notably the swelling of Kwon et al.^5^ is only partially attenuated by TTX, indicating that as in our experiments the structural change does not require action potentials. Their changes were at the nanometre scale, below the resolution of our diameter measurements, so we cannot rule out additional nanometre-scale plasticity of the node itself. While we found that axon plasticity occurred independently of calcium currents, we cannot rule out calcium from intracellular sources contributing, consistent with the calcium and calpain dependent axon plasticity previously reported^3^.

The problem of how a subthreshold stimulus engages Kv1 channels is resolved by the local electrochemistry of stimulation. Cathodal current alkalinizes the nearby extracellular space, and this alkalinization, coincident with a subthreshold depolarization, is sufficient to trigger axon thinning. This alkaline shift is local to the cathodal electrode tip, and easily missed without a pH sensitive microelectrode placed next to that tip. A local alkaline shift may increase K⁺ efflux through a pH-sensitivity of K2P or Kv channels^45^, raising periaxonal K⁺ and easing Kv1 activation so that the K⁺ build-up grows regeneratively. We cannot say which channel transduces the pH shift, and a K2P block does not settle it, since removing a resting leak depolarizes the axon and itself eases Kv1 activation, though involvement of Kv1 is established by its block by 4-AP. We also cannot rule out direct effects of pH on the bilipid membrane, since extracellular alkalinization reduces proton screening of fixed headgroup charges and can alter bilayer curvature or generate bilayer couple stress^46,47^. Overall, these results indicate that axon plasticity from subthreshold spinal cord stimulation is augmented by a coincident pH shift and depolarization. In contrast, suprathreshold stimulation, with intense spiking that readily activates Kv1.2 currents, likely produces plasticity without a coincident pH shift.

This pH action also explains why the same plasticity is produced by axonal GABA_A_ receptors, which simultaneously depolarize the axon^16^ and alkalinize the extracellular space through bicarbonate efflux^31–33^. Bath GABA or muscimol reproduce the full syndrome of axon shrinking and occlude the response to stimulation, identifying endogenous nodal GABA, supplied physiologically by spinal GABAergic neurons and astrocytes onto these axons^16^, as a natural trigger of the structural plasticity that we uncovered with an imposed current. These GABAergic neurons are themselves driven by afferent input, so that natural or stimulus-evoked activity in the sensory axons feeds back onto their own nodes^16,18^ to enhance plasticity. We had not anticipated that glycine receptors were also involved, with GABA and glycine application together more effective than GABA alone in shrinking axons. Glycine receptors are expressed by sensory afferents^19^, and likely act similarly to GABA_A_ receptors, though future studies are needed to test this. We used strong enough electrical stimuli to induce plasticity without endogenous GABA or glycine, since the observed axon plasticity dV is resistant to gabazine and strychnine. However, weaker stimulation likely relies on endogenous tonic GABA receptor activity on nodes^16^ derived from astrocytes or neurons for axon plasticity, because the increased excitability from such stimuli is blocked by GABA_A_ receptor antagonists or astrocyte toxins in vivo^7,48^, and is partly weakened by persistent sodium current blockers^49^.

The same process underlying axon plasticity with physiological activation can become pathological. Where axons are mechanically weakened, GABA and glycine drive not uniform thinning but localized swelling and frank beading, the precursor to axotomy and spheroid formation^42,43^. This may matter after concussion, where axons are mechanically strained without being severed. A second hit may then come not from further trauma but from ordinary GABAergic activity, and general anaesthetics that potentiate GABA_A_ receptors^50^ may carry particular risk in that window^51,52^. Activity-dependent osmotic remodelling thus sits on a continuum between physiological tuning and injury, and conditions that combine raised activity, altered potassium buffering or compromised myelin with mechanical vulnerability, may push it past the reversible regime, such as in demyelinating disease, traumatic brain injury, neuromyelitis and ischaemia^53,54^.

Functionally, activity-dependent axon thinning offers a slow, lasting means of setting the excitability and conduction of central myelinated axons. Because the node and paranode are spared while the internode thins, the change lowers the threshold for activating the axon and may help GABA secure conduction through branch points, where central afferents are most prone to failure and where nodal GABA supports propagation and proprioception^16^, though as detailed below this action can be complex. Engaged physiologically by GABA and by patterned activity, this plasticity is a candidate cellular substrate for the long-lasting changes in sensory and motor function that follow altered afferent activity^6,7^, giving a structural account of how an axon can carry a memory of its recent activity.

Every route to thinning that we have studied here, whether electrical, pharmacological or optogenetic, acts on many axons at once, as does the physiological GABAergic input to these axons, and the periaxonal K^+^ that escapes each axon, which enters a nodal extracellular space shared with its neighbours. Thus, the K^+^ build-up may be cooperative, with co-activated axons sustaining each other’s periaxonal K^+^ and overwhelming the glial K^+^ handling. This is supported by our finding that intracellular depolarizations of single axons failed to produce any sign of sustained axon thinning.

Activity-dependent internode thinning may underlie some of the clinical benefits of epidural spinal stimulation, which produces lasting improvements in sensory and motor function with stimuli often too weak to evoke action potentials^6,10,12,55–59^, consistent with the structural change we detail here for both short pulses and high frequency stimulation mimicking that used clinically. This may occur even for charge-balanced stimulation, as the electrochemical cathodal reactions that aid the axon plasticity can be irreversible. Slight charge imbalances amplify this^30^, especially as the currents used clinically are up to a thousand times higher than the weakest 1 μA currents that we find induce axon plasticity, and we found that both charge-balanced and kHz protocols indeed produced evidence of axon thinning. Even perfectly charge-balanced biphasic pulses at 20 μA and about 100 Hz, close to the parameters we used, produce irreversible hydrogen evolution at platinum microelectrodes once the charge density exceeds about 0.3 mC cm^−2^, ^60^ and pit platinum electrodes in vivo at 0.25 mC cm^−2^ per phase^61^, with associated local pH shifts^62^, and our 30 μm platinum-iridium wires likely approach this point at the charge densities we found activated axons (30 μA × 0.2 ms / 7.1 × 10^−6^ cm^2^ tip area gives 0.85 mC cm^−2^, before any correction for surface roughness). Suprathreshold high frequency firing from propagated spikes, without any electrochemical reaction, also drives the same thinning^4^, so stimulation that recruits spikes engages the mechanism as well, and both regimes may act through one structural plasticity. The subtle electrochemical pH effects of spinal cord stimulation are only needed to bring Kv1 channels to threshold to trigger axon plasticity during subthreshold stimulation, whereas suprathreshold stimulation directly activates Kv1 channels during spiking, likely inducing axon plasticity without the need for electrochemical actions, thus generalizing the phenomenon. Even transcutaneous spinal cord stimulation^56^ might engage this mechanism, since skin and dorsal column stimulation strongly activates GABAergic neurons that provide GABA to sensory axons^16^, which we show here triggers axon shrinking.

Future studies are required to follow up on a number of clinically relevant open questions raised that we did not pursue. For example, why did the CAP sometimes fail with strong stimulation? While moderate stimulation leads to axon internode shrinking that lowers the threshold to activate a node, stronger stimulation may cause excessive shrinking that shortens the axon length constant so much that conduction is slowed and eventually fails outright. Also, why did we only see internode shrinking, while previous studies have shown internode shrinking combined with nodal swelling^4^? Perhaps, the longer and suprathreshold stimulation used in the latter overwhelms the physiological adaptation process, again leading to loss of sensory transmission. These issues then bear on the question of how epidural or spinal cord stimulation functions, with activity-dependent axon plasticity providing a possible spectrum ranging from improved conduction to blocked conduction, depending on the intensity and duration of the stimulation, recruitment of GABAergic neurons, and desired outcomes. Ultimately, this may help explain the wide range of effective spinal cord and brain stimulation methods used clinically^35,63–65^, on one hand weak, possibly subthreshold, stimulation decreasing natural sensory transmission locally at the stimulation level, and thus reducing spasticity and pain, and on the other hand stronger suprathreshold stimulation increasing sensory transmission and proprioception more globally by the afferent-driven neuronal circuits activating GABAergic neurons. In summary, while our results leave many open clinical and physiological questions, including limitations in understanding the poorly accessible periaxonal space, they open up a new avenue to understand activity-dependent axonal plasticity and improve the clinical benefits of spinal cord or nerve stimulation.

## METHODS

### Animals and ethics

Recordings were made from myelinated dorsal column axons and motor axons in the sacrocaudal spinal cord of adult mice (3 to 6 months old), with male and female animals used in equal numbers. The adult sacrocaudal cord was chosen because it survives whole in vitro, unlike the lumbar cord, leaving the axonal architecture intact. Effects in male and female animals were similar and were grouped together. All procedures were approved by the University of Alberta Animal Care and Use Committee, Health Sciences division, and conformed to the guidelines of the Canadian Council on Animal Care. Most experiments used C57BL/6J mice (Jackson Laboratory). For optogenetic activation of GABAergic neurons we used GAD2//ChR2 mice, bred and induced with tamoxifen as previously detailed^16^. Briefly, heterozygous Gad2^CreER^ mice (Jackson Laboratory, 010702) were crossed with homozygous R26^LSL-ChR2-EYFP^ mice (Jackson Laboratory, 012569), and Cre was induced in adult mice with two doses of tamoxifen (0.2 mg/g, i.p.) at 4 to 6 weeks old, with recordings made more than 1 month later.

### In vitro preparation of the whole adult spinal cord

Mice were anaesthetized with urethane (0.11 g per 100 g, to a maximum of 0.065 g), a laminectomy was performed, and the entire sacrocaudal spinal cord was rapidly removed and immersed in oxygenated modified artificial cerebrospinal fluid (mACSF), as detailed previously^16^. The animal was then euthanized with Euthanyl (BimedaMTC; 700 mg per kg). The dorsal and ventral roots were left attached at the sacral S2, S3 and S4 and caudal Ca1 segments on both sides. After 1.5 h in the dissection chamber at 20°C, the cord was transferred to a recording chamber perfused with normal ACSF (nACSF) at 23°C and a flow rate above 3 mL per min. A 1 h wash in nACSF preceded recording, after which the nACSF was recycled in a closed system. The cord was secured to the Sylgard floor of the chamber with insect pins through connective tissue and cut root fragments, with its dorsal surface uppermost for stimulating and recording dorsal column axons and dorsal roots.

For recording from axons we mounted the freshly cut dorsal roots onto Ag/AgCl wires just above the bath and covered them in grease (a 3:1 mixture of petroleum jelly and mineral oil, with 9% wt silicon carbide powder) over about a 5 mm length (longer length provides an improved seal, as detailed below). The grease was applied on the roots as close as possible to the spinal cord, maximizing the recorded signal. Recording return and ground wires were likewise Ag/AgCl, and connected to the bath by a saline bridge. This bridge was made from saline and 4% agar in a 1 cc syringe, which we found essential to prevent bath-applied drugs like gabazine from contacting the wires and changing the junction potential, which can for gabazine be large enough to cancel the small recorded axon signal.

### Drugs and solutions

Two artificial cerebrospinal fluids were used: a modified ACSF (mACSF) in the dissection chamber and a normal ACSF (nACSF) in the recording chamber. The mACSF contained (in mM) 118 NaCl, 24 NaHCO_3_, 1.5 CaCl_2_, 3 KCl, 5 MgCl_2_, 1.4 NaH_2_PO_4_, 1.3 MgSO_4_, 25 D-glucose and 1 kynurenic acid. The nACSF contained (in mM) 122 NaCl, 24 NaHCO_3_, 2.5 CaCl_2_, 3 KCl, 1 MgCl_2_ and 12 D-glucose. Both were saturated with 95% O_2_ and 5% CO_2_ and held at pH 7.4. A HEPES-buffered nACSF was sometimes used and contained (in mM) 122 NaCl, 25 HEPES, 2.5 CaCl_2_, 3 KCl, 1 MgCl_2_ and 12 D-glucose. The solution was saturated with 100% O_2_ and adjusted to pH 7.4. An isotonic KCl solution was also used in which all NaCl was replaced by KCl with a final osmolarity the same as normal ACSF. Drugs added to the nACSF are summarized in Supplementary Table 1, with their targets, and NaCl was sometimes reduced when a drug had a concentration over a few mM to maintain equal osmolarity. Na^+^ and mixed cation channels were blocked with TTX (TTX-citrate; 2 - 50 μM, NaV), riluzole (50 μM, persistent NaP), ZD7288 (50 μM, HCN), ruthenium red (10 μM, TRP, tested in HEPES), flufenamic acid (FFA; 300 μM, TRP, tested in HEPES), phenamil mesylate (300 μM, ASIC and ENaC, tested in HEPES), and benzamil (100 μM, ENaC and NCX, tested in HEPES). Ca^2+^ channels and stores were targeted with cadmium chloride (400 μM, CaV), nimodipine (20 μM, CaV1.3 L-type), ZnCl_2_ (3 mM, CaV3 T-type), dantrolene sodium salt (50 μM, ryanodine receptors), cyclosporin A (10 μM, mitochondrial permeability transition pore and calcineurin), and Ca^2+^-free medium with EGTA (5 mM, tested in HEPES). K^+^ channels were blocked with 4-aminopyridine (4-AP; Kv1) at low (100 μM) and high doses (1 to 3 mM), divalent cations (CSF with 15 mM MgCl_2_ and 6.5 mM CaCl_2_ with NaCl reduced to match osmolarity, Kv), XE991 (50 μM, Kv7), citalopram (500 μM, K2P), quinidine (100 μM, K2P, TASK and Ca-activated K, tested in HEPES), tertiapin (0.5 μM, Kir), Ba^2+^ chloride (10 mM, Kir), and isotonic KCl (to collapse E_K_). Cl^−^ and anion channels and transporters were targeted with: gluconate^−^ (replacing Cl^−^ in nACSF at equal molarity), tamoxifen (100 μM, VRAC), ZnCl_2_ (3 mM, CLC), DIDS (0.5 mM, anion exchanger, NBC and VRAC, tested in HEPES), S0859 (200 μM, NBC), EIPA (100 μM, NHE1, tested in HEPES), bumetanide (30 μM, NKCC1, tested in HEPES), VU0463271 (1 μM, KCC2), DIOA (20 μM, KCC2), and TGN020 (300 μM, AQP4).

The Na/K-ATPase was blocked with ouabain (low dose 100 μM, high dose 1 mM) or by replacing bath Na^+^ with N-methyl-D-glucamine^+^ (NMDG^+^; 122 mM NMDG^+^ in HEPES-buffered nACSF with pH adjusted to 7.4 with HCl). Receptors were blocked with CNQX and APV (50 μM, glutamate), gabazine or bicuculline (50 μM, GABA_A_), strychnine (5 μM, glycine), CGP55845 (1 μM, GABA_B_), tubocurarine hydrochloride (30 μM, ACh), brilliant blue (30 μM, P2X), and suramin (100 μM, P2). Carbenoxolone disodium (100 μM) was used to block gap junctions (connexins), H89 dihydrochloride (20 μM) to block PKA, and L-AAA (1 mM) as an astrocyte toxin. Acetazolamide (100 μM) with HEPES-buffered ACSF was used to reduce bicarbonate. GABA (2 mM), glycine (2 mM, glycine receptors), and muscimol (15 μM, GABA_A_) were applied to activate axonal receptors. The HEPES-buffered nACSF alone did not change the results, in control trials. Drugs were obtained from Tocris, Sigma-Aldrich and Toronto Research Chemicals, prepared as 10 to 50 mM stocks in water or DMSO and diluted to final concentration in nACSF, with the final DMSO concentration kept below 0.04%, which by itself had no effect on the recorded axon signals in vehicle controls. Local pH was raised where indicated by brief application of NaOH.

### Focal subthreshold stimulation

Focal cathodal stimulation was delivered to the surface of the dorsal columns through a large diameter blunt tungsten electrode (tip of about 55 μm, 15 kΩ impedance, to minimize current density; TM53CCINS-SB, WPI), with a return Ag/AgCl anode wire in a 4% agar saline bridge connected to the bath. Cathodal current was passed from a constant-current stimulator (Isoflex, Israel) as a 1 to 3 s pulse at intensities below those that evoked an axonal volley, so that the stimulus was subthreshold for firing. In some experiments a saline bridge was interposed between the electrode and the cord, in which the electrode sat in a large HEPES-buffered reservoir (500 ml) that diluted any local pH shift and connected to a micropipette that delivered the 3 s current to the axons, separating the depolarizing action of the stimulus from its local electrochemistry. The same tungsten electrode on the dorsal columns was also used to evoke test action potentials in group I afferents with a 0.1 ms supra-threshold current pulse of 1.25 to 1.5 times threshold (T), which were recorded on the dorsal roots as a compound action potential (CAP).

### Local pH measurement

Extracellular pH near the stimulation site was measured with an 8 μm tip pH-sensitive microelectrode (pH-10, Unisense, Aarhus, Denmark) placed within about 200 μm of the tungsten electrode. The signal was referenced to the bath and calibrated against standard buffers spanning pH 6 to 9, allowing the local alkaline shift produced by cathodal current to be followed in time.

### Viral labelling of sensory afferents

Large diameter peripheral afferents were labelled by viral vector injections, as previously detailed^16^. Adeno-associated virus (AAV) vectors with the transgene encoding the cytoplasmic fluorophore GFP or tdTom under the CAG promoter were injected intraperitoneally into anesthetized P1 to P2 mice (AAV9-CAG-tdTom or AAV9-CAG-GFP, 5.9 × 10^12^ vg/ml, 2 to 4 μl per injection; UNC Vector Core).

### Optogenetic activation of GABAergic neurons

Light was delivered from a 447 nm laser (Laserglow Technologies, Toronto) through a fibre optic cable and a half cylindrical prism, giving a narrow beam focused on the dorsal spinal cord^16^. Silicon carbide powder (9% wt) was added to the grease on the roots to make it opaque, preventing light artifacts in the recording wire.

### Two-photon imaging of live axons

Live dorsal column axons labelled with AAV9-GFP or tdTom were imaged in the whole in vitro or in vivo spinal cord with an FV1000 MPE multiphoton scanning microscope (Olympus, Tokyo, Japan), using a MaiTai Ti:sapphire femtosecond pulsed laser at 810 nm excitation and an Olympus 20× immersion objective (NA 1.0). Axons were imaged near the stimulating electrode in z stacks within 50 μm of the dorsal surface, with 0.5 - 1 μm optical sections, up to 3 images averaged per section, and stacks repeated continuously to follow axon changes over time. Mean axon diameter was measured adjacent to the stimulation zone with one internode distance^16^. We focused on large group I and II sized axons, though all axons behaved similarly, shrinking with stimulation. Net glia size was estimated from the gap between adjacent axons as described in the Fig 1 legend.

### Immunohistochemistry and confocal imaging

After stimulation (∼1 h after) in vitro spinal cords and adjacent dorsal roots were dropped in 2 or 4% paraformaldehyde for 2 h, and then cryoprotected in 30% sucrose in phosphate buffer for about 48 h. For this only the left side of the spinal cord was stimulated and the right side served as control non-stimulated tissue, in matched pairs rostro-caudally used to perform pairwise comparisons of stimulation effects. Cords were then embedded in OCT (Sakura Finetek), frozen at −60°C with 2-methylbutane, cut on a cryostat (NX70, Fisher Scientific) in sagittal or transverse 25-μm sections, and mounted on slides.

Sections were rinsed with phosphate-buffered saline (PBS, 100 mM) and then with PBS containing 0.3% Triton X-100 (PBS-TX). Nonspecific binding was blocked with a 1 h incubation in PBS-TX with 10% normal goat serum (NGS; S-1000, Vector Laboratories) or normal donkey serum (NDS; ab7475, Abcam). Sections were then incubated for at least 20 hours at room temperature with a combination of the following primary antibodies in PBS-TX with 2% NGS or NDS: rabbit anti-Caspr (1:500; ab34151, Abcam, Cambridge, UK), mouse anti-Caspr (1:500; K65/35, NeuroMab, Davis, USA), rabbit anti-GFP (1:500; A11122, ThermoFisher Scientific, Waltham, USA), rabbit anti-RFP (1:500; PM005, MBL International, Woburn, USA), mouse anti-Pan Sodium Channel (1:500; S8809, Sigma-Aldrich, St. Louis, USA), mouse anti-Kv1.2 (1:500; L76/36, NeuroMab, Davis, USA), guinea pig anti-GFAP (1:500; 173 308, Synaptic Systems), and mouse anti-CNPase (1:200; MAB326, Millipore). Genetically expressed GFP and tdTom were amplified with the anti-GFP and anti-RFP antibodies, rather than relying on endogenous fluorescence.

The following day, sections were rinsed with PBS-TX and incubated for 2 h at room temperature with the appropriate fluorescent secondary antibodies (Alexa Fluor conjugates, 1:200 to 1:500, raised in goat or donkey; Thermo Fisher Scientific and Abcam) in PBS-TX with 2% NGS or NDS. When anti-mouse antibodies were applied to mouse tissue, the Mouse on Mouse immunodetection kit (M.O.M; BMK-2201, Vector Laboratories) was used before applying antibodies, including a 1 h incubation with a mouse Ig-blocking reagent. After rinsing with PBS-TX and PBS, slides were coverslipped with Fluoromount-G (00-4958-02, Thermo Fisher Scientific).

Standard negative controls, in which the primary antibody was either omitted or blocked with its antigen, confirmed the selectivity of the staining, and no specific staining was seen in these controls.

Images were acquired by confocal microscopy (Leica TCS SP8) with a ×63 (1.4 NA) oil immersion or ×20 water immersion objective, taking 0.1-μm optical sections collected into a z-stack over 10 to 20 μm. Excitation and recording wavelengths were set to optimize the selectivity of imaging the fluorescent secondary antibodies, and large areas were imaged with the Tilescan option in Leica Application Suite X (Leica Microsystems).

Axon node, paranode and juxtaparanode diameters were measured for all axons in each image of control and stimulated dorsal columns. These were sorted by node diameter and the top 50% were analysed, which assured that we sampled a similar population of axons with and without stimulation, roughly corresponding to the group I and II sized axons that make up about half the myelinated fibres in these roots^66^. Nodal gap ratio was also computed as the ratio of the node length over diameter.

### Data analysis and statistics

Recordings were acquired in Axoscope and analysed in Clampfit 10 (Axon Instruments and Molecular Devices), Excel (Microsoft) and SigmaPlot 14.5 (Systat Software). The grease-gap DC potential was amplified, low-pass filtered at 10 kHz and sampled at 30 kHz (Axoscope; Axon Instruments and Molecular Devices). The shrinkage-related depolarization dV was quantified 10 to 20 min after stimulation, when the early active currents had subsided. Dorsal roots and associated axons were recorded and stimulated at widely separated segmental locations within the spinal cord and thus were treated as independent, so when testing drug actions on axon plasticity with stimulation statistics were computed across all roots (n) from all animals, with 3 to 5 animals per drug (as detailed for dV in Supplementary Table 1). The overall effects of axon stimulation on axon plasticity from all animals prior to adding drugs were computed from means across animals (n, in main text). A Student t test or one-way ANOVA, as appropriate, was used to test for statistical differences between two or more independent comparisons respectively, with a significance level of p < 0.05, all two-sided. Tests were paired for pre-treatment and post-treatment data but otherwise were unpaired. Multiple comparisons with t tests or ANOVA were followed by a post hoc Bonferroni correction or Tukey test to determine which pairs of measures likely differed while compensating for the multiple comparisons indicated. A Wilcoxon signed-rank test was used where data were not normally distributed. Data are given as mean ± standard deviation (SD). Power analysis was used a priori to design experiments and determine sample sizes n (with α = 0.05, β = 0.2, r = 0.8 to 1.6, t test or ANOVA, where r = effect size, based on previous studies). Animals were randomly allocated to different groups for the in vitro and in vivo experiments using a block design. Data collection and analysis were not performed blinded to the conditions of the experiments, but the collection and analysis was automated so that the experimenter had no influence on the outcome.

### Grease-gap recording of the membrane potential and internodal axial resistance

Composite axonal potentials were recorded by grease gap from the proximal end of a cut dorsal root, with the recorded potential reflecting the transmembrane potential Vm of the nodes in the bath adjacent to the grease seal, as detailed previously^16^. The grease-gap electrode records the transmembrane potential of the axons in the root, sensed as a voltage V across the extracellular resistance of the grease seal. The recorded potential V is strictly proportional to the axon membrane potential.

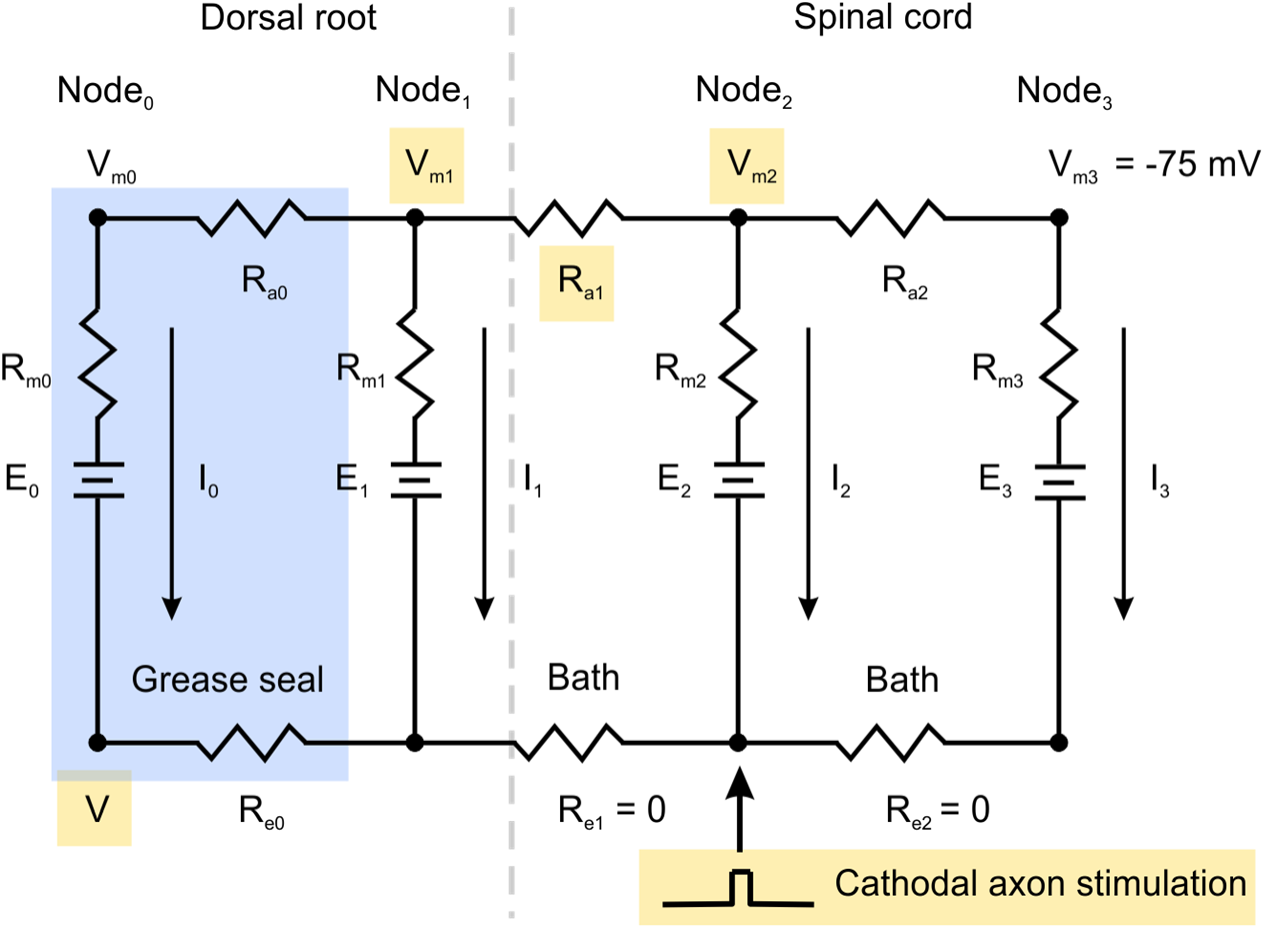

*Resistive ladder representing axon used to derive equations (2) and (8). Rung currents I_0_ to I_3_ pass down each node membrane and battery E. In steady state the spinal cord provides a steady axial current across R_a1_, making the recorded potential V sensitive to this axonal resistance*.

To see this we modeled the root as a ladder of nodes, each a rung of membrane resistance Rm in series with a potassium battery E, joined along the axoplasm by axial resistances Ra and along the outside by extracellular resistances Re. Assuming that node_0_ lies in the grease and the remaining nodes lie in the bath, then the recorded potential V is the voltage drop across its seal resistance R_e0_. To relate this to the membrane potential, consider node_0_ together with the adjacent node_1_ in the bath that has a membrane potential V_m1_. Summing the voltage drops around the loop between these two nodes we get (by Kirchhoff’s law): V_m1_ − E_0_ = I_0_ R_a0_ + I_0_ R_m0_ + I_0_ R_e0_, where E_0_ is the fixed reversal potential (battery) of the node in grease. Then combining with the voltage drop across the seal, V = I_0_ R_e0_, we get the voltage divider equation of the grease gap:

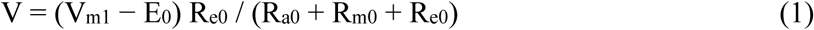

where E_0_ is a fixed voltage. Several simple observations follow from this equation. 1) When the grease seal is poor R_e_ approaches zero and V gets very small and not useful. 2) When the seal is nearly perfect with R_e_ very large then V = V_m1_ − E_0_ and so except for the fixed offset E_0_ the recorded signal is exactly the axon potential. In our experience in large ensheathed roots this situation is difficult to achieve even with using sucrose rather than oil as an insulator, because the fluid between the axons is difficult to displace. 3) With a good grease seal we find that R_e0_ is about equal to R_a0_ + R_m0_ so V = 0.5 (V_m1_ − E_0_), as detailed below, with the recording still accurately reporting V_m_, but with a fixed attenuation. 4) When we measure changes in potential dV and dV_m_ caused by stimulation or drug additions then differencing equation (1) gives the key grease-gap relationship:

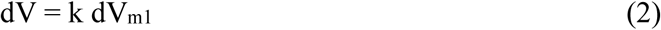

Here k = R_e0_/(R_a0_ + R_m0_ + R_e0_). The fraction k is set by the node_0_ resistances in the grease and does not change when a distant internode shrinks. We estimated k by collapsing V_m1_ to zero in isotonic KCl, which gave a change in V of 32 mV, so that with V_m1_ resting at about −65 mV, as shown below from equation (8), k = 32/65 ≈ 0.5.

We next relate V_m1_ to the axial resistance of the shrinking internode. Writing the Kirchhoff voltage and current laws around each successive loop for the four nodes, with currents I_0_, I_1_ and I_2_ through rungs 0 to 2 and each rail segment carrying the running sum of upstream rung currents, gives

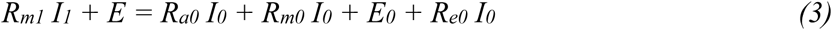

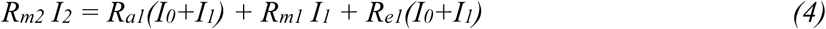

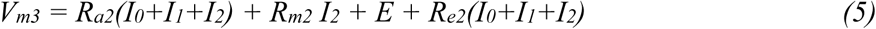

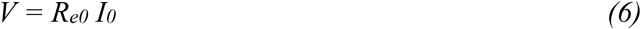

where V is the recorded potential, developed across the grease seal resistance R_e0_, V_m3_ is the membrane potential of node_3_, and E is the fixed reversal potential of nodes in the bath (near −85; E = E_1_ = E_2_ = E_3_). For simplicity, to start we compute the solution without node_3_, by taking R_a2_ to be large, so that negligible axial current passes beyond node_2_, and node_3_ does not enter the derivation (though similar conclusions hold without this assumption, as detailed later). Intracellular recordings from these axons in the dorsal columns shows that they rest at −75 mV while the cut dorsal roots are in grease^16^. Thus assume V_m2_ is fixed at −75 mV. Also assume we stimulate at node_2_ and the axial resistance R_a1_ joining it to node_1_ in the root is what changes. The axon beyond the grease lies in the bath, which shorts the extracellular rail between the intact nodes, so R_e1_ = R_e2_ = 0 and the recorded potential is the drop across the grease seal alone (equation (6)), V = R_e0_ I_0_, which by the voltage divider of equation (1) is V = (V_m1_ − E_0_) R_e0_/(R_a0_ + R_m0_ + R_e0_), depending only on V_m1_ and the fixed offset E_0_.

The axial voltage drop along the axoplasm between node_1_ and node_2_ gives

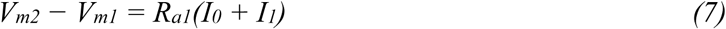

Now I_0_ + I_1_ = − I_2_ and V_m2_ = R_m2_ I_2_ + E, so R_m2_(I_0_ + I_1_) = −(V_m2_ − E). Substituting into equation (7) gives

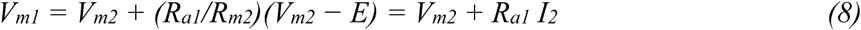

Equation (8) implies that the recorded node in the root is always more depolarized than the axon in the spinal cord. Specifically, the recorded node_1_ in the root is depolarized (V_m1_) because it sits at the sealed end of a short cable carrying axial current, and so has to pass the entire axial current rather than pass some onward to another node, a standard boundary effect that generalizes to many nodes and not just the few discussed here^17^. The spinal cord acts as a large reservoir current sink that holds node_2_ at approximately a fixed potential, and the node in grease acts as a source, depolarized relative to the nodes in the bath as expected at a cut end sealed by grease, leading to a standing axial current flowing through the internode resistance R_a1_ which causes a voltage drop that depolarizes node_1_. As the internode narrows its axial resistance rises, so the drop and the recorded depolarization grow, giving a direct readout of shrinkage. For these axons the internodal axial resistance is comparable to the leak-dominated nodal membrane resistance, R_a1_ ≈ R_m2_, both of order tens of MΩ^67,68^, so equation (8) places the resting potential of node_1_ at V_m1_ ≈ −65 mV, when as usual V_m2_ = −75 mV. Together with the isotonic KCl measurements, this implies that V_m1_ rests at −65 mV and E_0_ is near zero, so that V = kV_m1_.

Combining equation (8) with the grease-gap voltage divider law (equation (1)) V = k (V_m1_ − E_0_) with k = R_e0_/(R_a0_ + R_m0_ + R_e0_) gives the key relationship between the recorded potential and the internodal axial resistance,

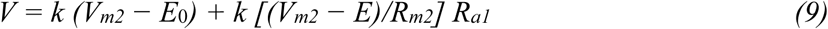

Because E_0_, E, R_m2_ and V_m2_ are assumed to be relatively fixed in the steady state before and after stimulation, the recorded potential varies linearly with the axial resistance R_a1_. Differentiating equation (9) gives the recorded potential change dV when the axial resistance changes,

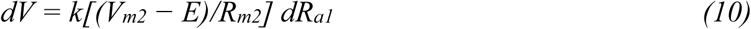

The full solution to equations (3) to (6), without assuming R_a2_ is large or V_m2_ fixed, gives the same resting values, V_m1_ ≈ −64 mV and V ≈ −32 mV, and the same linear dependence of V on R_a1_ as equation (9), though with a slope about half that of equation (10), since V_m2_ is not perfectly clamped. Equation (10) therefore sets an upper bound on the recorded signal for a given shrinkage.

The axial resistance of a cylindrical internode is set by its radius,

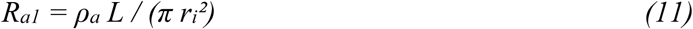

so that when the internode shrinks to a fraction f of its initial radius, the axial resistance rises by the ratio

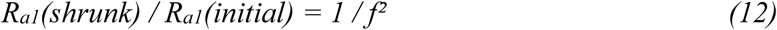

and so dR_a1_ = R_a1_(1/f² − 1). Substituting this into equation (10) gives the recorded change with axon shrinking:

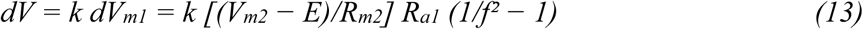

For a resting node at V_m2_ near −75 mV, the potassium reversal at E = −85 mV, and R_a1_ ≈ R_m2_ as above, equation (13, middle term) gives a true depolarization dV_m1_ of about 10 mV in a stimulated axon for shrinkage to 70% of the radius (f = 0.7). With an estimated V_m1_ of −65 mV for the axon in the root at rest, this dV_m1_ indicates that after stimulation V_m1_ was at about −55 mV. Diluted by the roughly one in 10 axons of a root that are stimulated, and read through the fixed voltage divider (k = 0.5), this corresponds to a recorded dV of about 0.5 mV, matching the observed signal. A recorded deflection therefore maps directly onto the fractional change in diameter, providing the quantitative link between the dV and the physical shrinkage.

We confirmed that dV resulted from an axial current and thus reports an axial-conductance change in steady state after active currents subside, by showing that after dV was induced by stimulation it was abolished when the axial current was cancelled by injected or induced current (significantly reduced dV to 0.80 ± 1.07 % of initial value of 100%, p < 0.05, n = 5). We did not routinely cancel the axial currents with current injection like this, or more generally voltage clamp the axons and measure current, because the large size of the roots required currents that often saturated our amplifier, exceeding its compliance.

### Dependence of the signal on the length of root in the grease

In preliminary trials we found that the recorded amplitude V is larger when more nerve length and nodes lie within the grease seal, and so we used about 5 mm of nerve in the grease. The reason for this is seen by the following analysis. We compare one node against two nodes lying within the grease seal, treating the root as a uniform ladder of passive axons governed by equations (3) to (6). Writing R_m_ for the membrane resistance of a node and W = R_a_ + R_e_ for the series resistance per section, one node in the grease gives the single-node divider of equation (1), here V = V_m_ R_e_/(R_m_ + W), while two nodes in the grease give V = V_m_ R_e_(3R_m_ + W)/(R_m_² + 3R_m_W + W²). Both recorded potentials are always smaller than the true nodal potential, since the extracellular resistance is only part of each loop. For a fixed nodal potential the two-node measurement always exceeds the one-node measurement, because their difference, V_m_ R_e_ R_m_(2R_m_ + W)/[(R_m_ + W)(R_m_² + 3R_m_W + W²)], stays positive, so including more nodes within the grease yields a larger recorded potential, accounting for the larger signals obtained with longer lengths of root in the grease. This gain saturates as the membrane resistance becomes large relative to the series resistance, where the two-node signal approaches three times the one-node value and the recorded potential approaches but never reaches the membrane potential; as the membrane resistance falls toward zero the rungs short the nodes together and additional nodes confer no benefit.

### Spatial localization of the dV signal to internodes near the grease

The grease-gap signal is weighted toward the internodes nearest the recording site. With the bath shorting the extracellular rail beyond the grease, a change in the axial resistance of a given internode reaches the electrode only after passing the intervening nodes, at each of which part of the current escapes through the nodal membrane to the bath. Solving the full four-node model of equations (3) to (6) for a change in the axial resistance R_a2_ of an internode two sections beyond the recorded node gives a recorded change

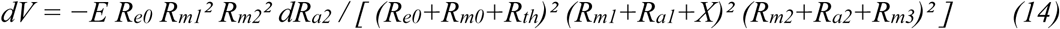

with X = R_m2_(R_a2_+R_m3_)/(R_m2_+R_a2_+R_m3_) and R_th_ = R_a0_ + R_m1_(R_a1_+X)/(R_m1_+R_a1_+X). The recorded change is attenuated by two squared current-divider factors, R_m1_²/(R_m1_+R_a1_+X)² and R_m2_²/(R_m2_+R_a2_+R_m3_)², each less than one. A change in the internode immediately adjacent to the recorded node carries a single such factor, an internode one section deeper carries two, and each additional section of separation adds another, so the contribution of an internode to the recorded signal falls off geometrically with its distance from the grease. The dV assay is therefore a local probe that preferentially reports shrinkage of the internodes closest to the cut end in the grease and progressively underweights shrinkage deeper in the bath. For this reason the stimulating electrode was positioned close to the recording site so that the shrinkage it evoked was faithfully reported, and absolute shrinkage of distant internodes is expected to be underestimated by the recording.

### Cable model of the excitability change produced by internodal thinning

To relate the measured internodal shrinkage to the observed change in excitability we used a cable model of the myelinated axon in which the membrane current is concentrated at the nodes of Ranvier and the internodes are treated as passive axial resistors with no transmembrane leak. We distinguish two radii: the node radius r_n_, which is spared and does not change, and the internode radius r_i_, which falls with shrinkage and sets the internodal axial resistance. For an internode of length L and axoplasmic resistivity ρ_a_, the axial resistance between adjacent nodes is R_a_ = ρ_a_ L / (π r_i_²), which scales as 1/r_i_². The nodal membrane terms depend only on the fixed node radius r_n_ through the nodal area 2π r_n_ ℓ_n_, where ℓ_n_ is the nodal length. Writing the current balance at node n, with membrane capacitance C_m_ and specific nodal membrane resistance R_m_ per unit area, gives the discrete nodal equation^15,69^

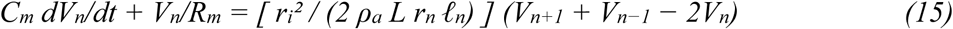

The coupling between nodes is carried entirely by the internodal term and scales as r_i_², since it is proportional to the axial conductance 1/R_a_, while the node radius r_n_ enters only the fixed membrane terms. Using a Taylor expansion of the terms in equation (15) one internode each way about the node gives V_n+1_ + V_n−1_ − 2V_n_ = L² ∂²V_m_/∂x² + (L⁴/12) ∂⁴V_m_/∂x⁴ + … For a near steady state exponential solution V ≈ A exp(−x/λ) (reached in a few μs, well within the 0.1 ms test pulse), with length constant λ, these terms are in the ratio (L/λ)^2^, (L/λ)^4^/12 and so on. Thus, when λ exceeds the internode length L, equation (15) reduces approximately to the continuous cable equation

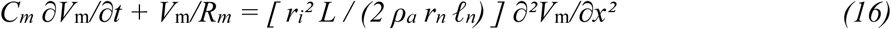

Here V_m_ = V for consistency with our earlier notation for membrane potential used above. The coefficient on the right is the only term that depends on the internode radius. Rescaling distance by the internode radius, X = x/r_i_, removes r_i_ from the equation entirely,

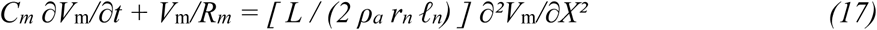

so the voltage profile over distance is universal in the rescaled coordinate and all of the radius dependence is absorbed into the coordinate stretch x = r_i_ X. Two simple scaling rules follow. First, the effective electrotonic length over which voltage spreads scales linearly with the internode radius, λ_eff_ ∝ r_i_, because the axial resistance scales as 1/r_i_² while the fixed node sets the membrane terms. Second, the current required to bring a node to threshold is the fixed per-area threshold drive times the membrane area over which it must be supplied, which is the node circumference 2π r_n_ times the spread length λ_eff_ ∝ r_i_, so

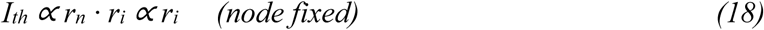

The threshold current therefore falls in direct proportion to the internode radius as the internode shrinks. For the measured shrinkage to about 70% of the initial diameter, the threshold current is predicted to fall to about 70% of its initial value, a substantial increase in excitability that arises purely from the geometry, with the node and its channels unchanged. This provides the quantitative cable-theoretic basis for the excitability increase that accompanies shrinkage. The same result follows directly from equation (15) without the continuous approximation. Letting x = Ln and seeking a steady state exponential V(x) = A exp(−x/λ) gives cosh(L/λ) = 1 + ρ_a_Lr_n_ℓ_n_/(R_m_r_i_^2^), so that for λ > L, λ ≈ r_i_ √(R_m_L/2ρ_a_r_n_ℓ_n_), again proportional to r_i_.

The reduced threshold also changes the compound action potential through the recruitment of additional axons. We assume the number recruited rises linearly with the applied current above threshold,

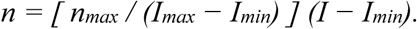

Expressing the current in multiples of the threshold current, J = I/T with I_min_ = T and J_max_ = I_max_/T, this becomes

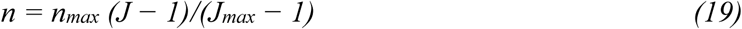

The compound action potential is proportional to n. Shrinkage lowers the threshold from T to fT, and so the ratio of equation (19) after and before shrinkage is

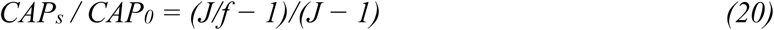

valid while J/f ≤ J_max_. Thus, for a stimulus applied near threshold, a modest reduction in threshold recruits a disproportionately large number of additional axons.

For example, if an axon shrinks by f = 0.7, then for the stimulus parameters that we used of 1.25 to 1.5 times threshold (J = 1.25 to 1.5) to study the fastest component of the CAP arising from group I afferents that are fully recruited at twice threshold (J_max_ = 2), the compound action potential increase is CAP_s_ / CAP_0_ = 2 to 3.1.

## ACKNOWLEDGMENTS

We thank Leo Sanelli and Jennifer Sauve for technical assistance. We thank Professor Elzbieta Jankowska for initiating the study, and many suggestions on the manuscript. We owe a great debt to our mentor and colleague RB Stein who motivated the analysis in this study and sadly passed away recently.

## DECLARATION OF GENERATIVE AI AND AI-ASSISTED TECHNOLOGIES

Claude was used to aid in language editing and to enhance readability during manuscript preparation. The authors carefully reviewed and revised the text and accept full responsibility for the published content.

## SUPPLEMENTARY TABLE

**Supplementary Table 1.**
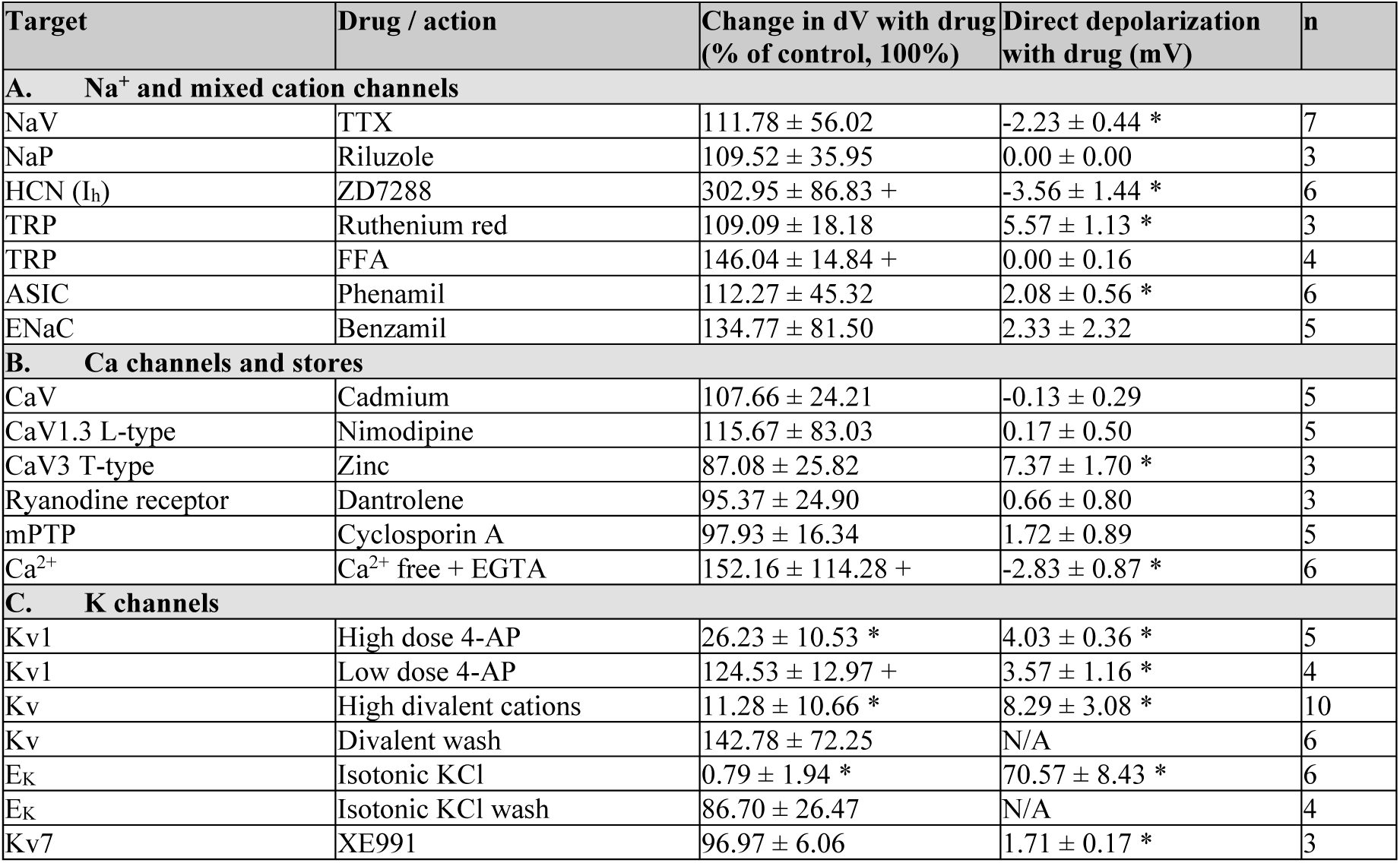
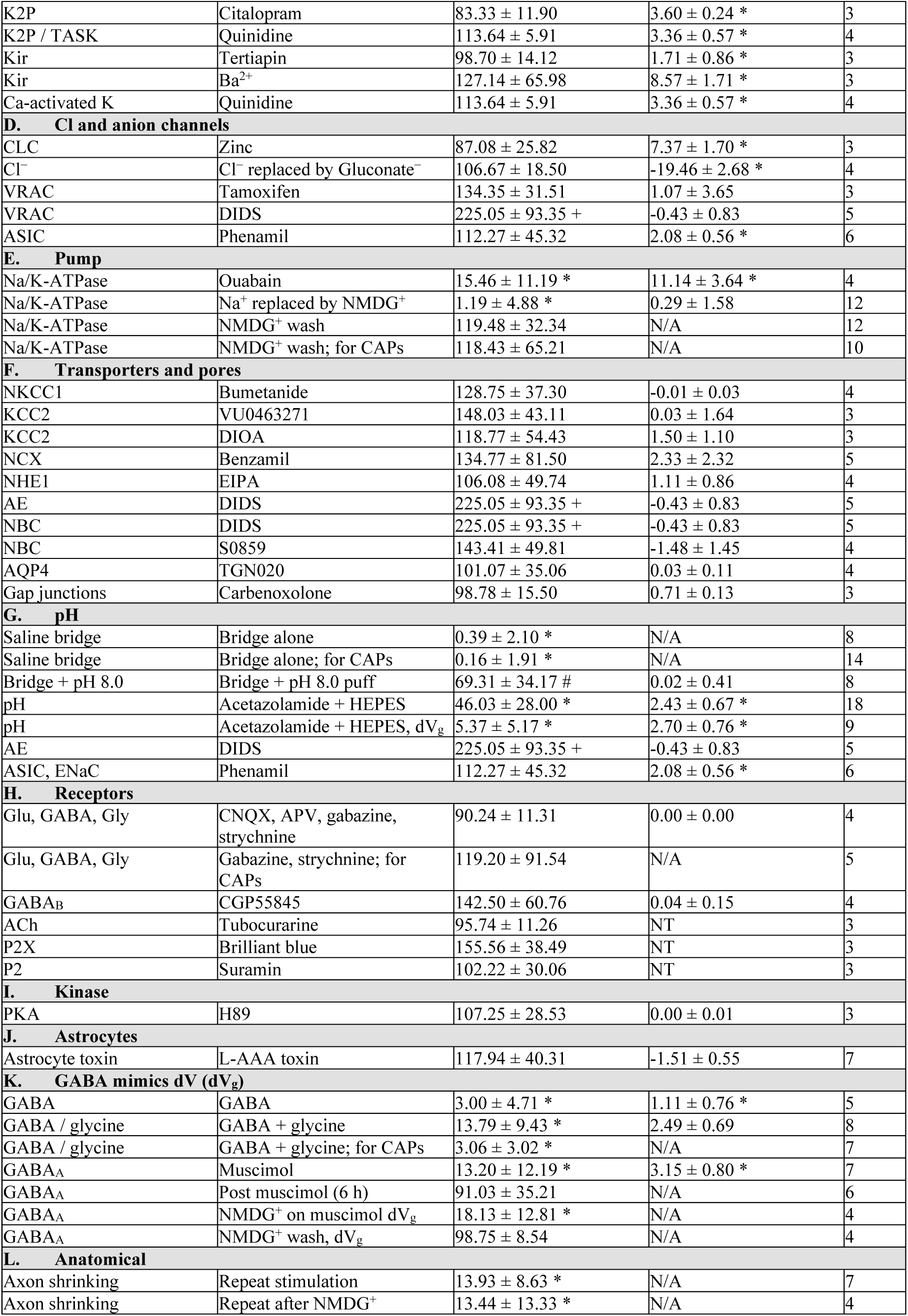
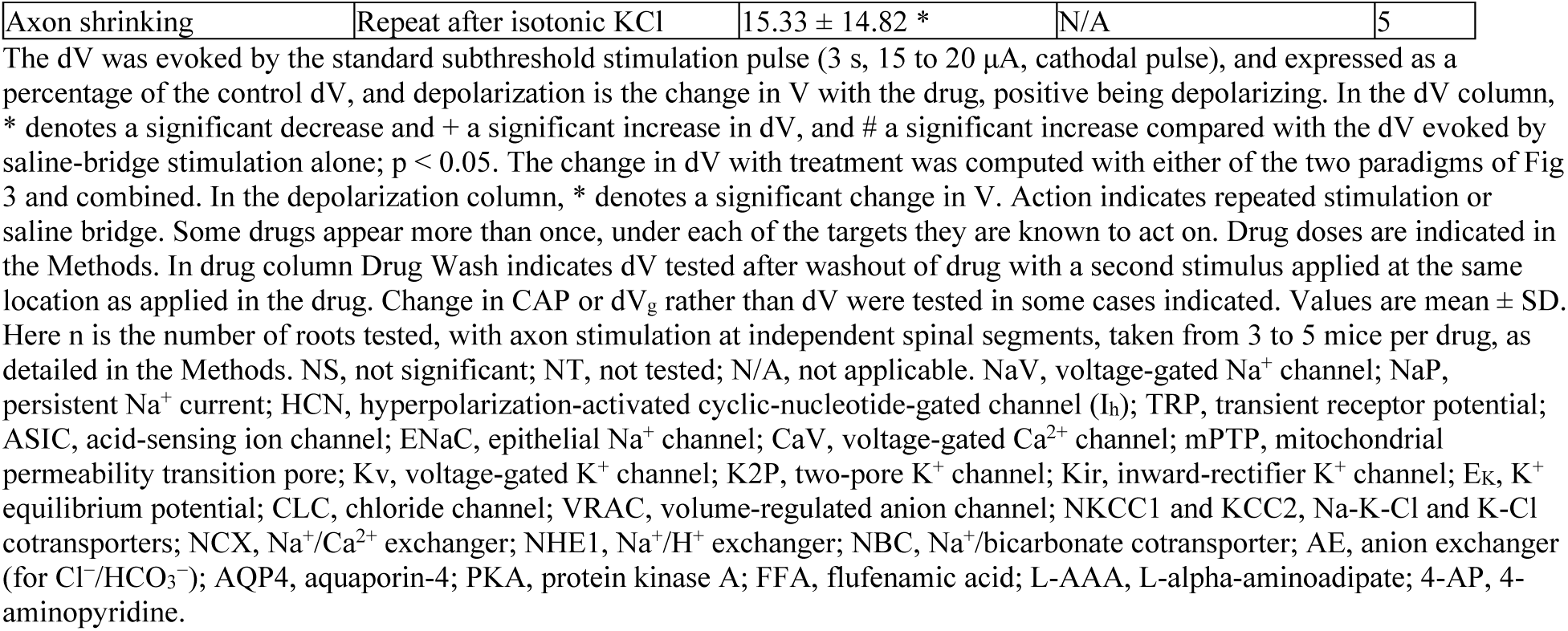
Effects of drugs and actions on the stimulus-evoked dV and resting membrane potential.

## SUPPLEMENTARY FIGURE

**Supplementary Figure 1.**
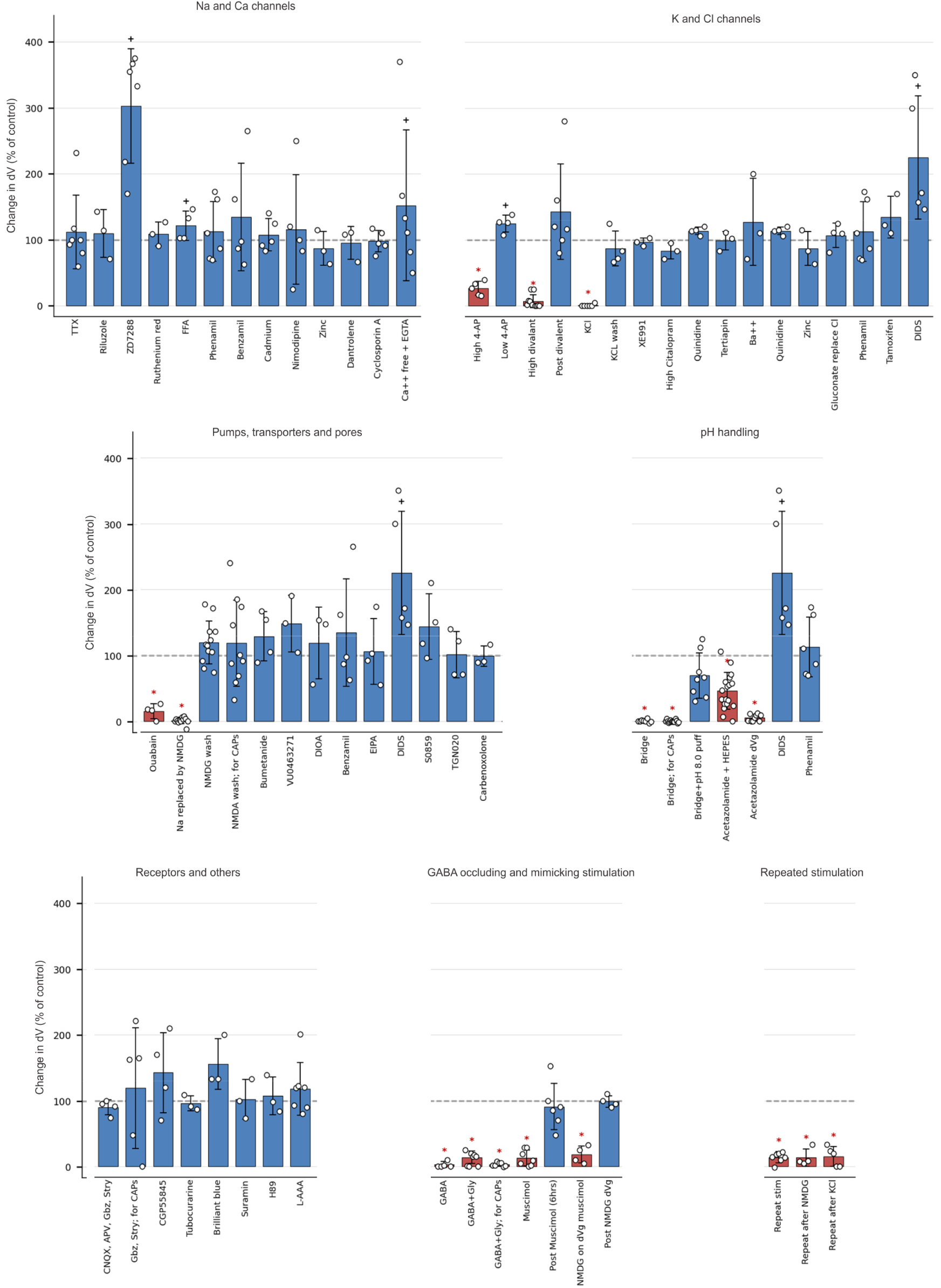
Effects of all drugs and actions on the stimulus-evoked dV. Bar graphs of the data summarized in Supplementary Table 1, grouped by target class, with the dV expressed as a percentage of the control dV evoked by the standard subthreshold stimulation (3 s, 15 to 20 μA, cathodal pulse). Dashed line, control at 100%. Circles, individual recordings. Bars, mean ± SD. Red bars and * denote a significant decrease in dV with the drug or action, and + a significant increase, p < 0.05. Group sizes and values are given in Supplementary Table 1.

